# Intestinal transit amplifying cells require METTL3 for growth factor signaling, KRAS expression, and cell survival

**DOI:** 10.1101/2023.04.06.535853

**Authors:** Charles H. Danan, Kaitlyn E. Naughton, Katharina E. Hayer, Sangeevan Vellappan, Emily A. McMillan, Yusen Zhou, Rina Matsuda, Shaneice K. Nettleford, Kay Katada, Louis R. Parham, Xianghui Ma, Afrah Chowdhury, Benjamin J. Wilkins, Premal Shah, Matthew D. Weitzman, Kathryn E. Hamilton

## Abstract

Intestinal epithelial transit amplifying cells are essential stem progenitors required for intestinal homeostasis, but their rapid proliferation renders them vulnerable to DNA damage from radiation and chemotherapy. Despite their critical roles in intestinal homeostasis and disease, few studies have described genes that are essential to transit amplifying cell function. We report that the RNA methyltransferase, METTL3, is required for survival of transit amplifying cells in the murine small intestine. Transit amplifying cell death after METTL3 deletion was associated with crypt and villus atrophy, loss of absorptive enterocytes, and uniform wasting and death in METTL3-depleted mice. Ribosome profiling and sequencing of methylated RNAs in enteroids and *in vivo* demonstrated decreased translation of hundreds of unique methylated transcripts after METTL3 deletion, particularly transcripts involved in growth factor signal transduction such as *Kras*. Further investigation confirmed a novel relationship between METTL3 and *Kras* methylation and protein levels *in vivo*. Our study identifies METTL3 as an essential factor supporting the homeostasis of small intestinal tissue via direct maintenance of transit amplifying cell survival. We highlight the crucial role of RNA modifications in regulating growth factor signaling in the intestine, with important implications for both homeostatic tissue renewal and epithelial regeneration.

## Introduction

The intestinal epithelium digests and absorbs nutrients, protects against pathogen invasion, and regulates interactions between mucosal immune cells and the gut lumen (1). These essential functions are made possible by the continuous renewal of differentiated intestinal epithelial cells (2). Epithelial renewal begins at the intestinal crypt base, where intestinal stem cells expressing the Wnt target gene, LGR5, initiate differentiation and migrate up the crypt wall. As they move up the crypt wall, LGR5+ stem cells differentiate into intestinal stem progenitors known as transit amplifying (TA) cells (2). TA cells rapidly undergo successive cycles of proliferation to generate the bulk of intestinal epithelium. While existing research emphasizes the role of LGR5+ stem cells in tissue renewal, TA cells are the primary site of intestinal epithelial proliferation and differentiation, and they produce the majority of differentiated epithelium (3, 4). Rapid proliferation is the central defining feature of TA cells, and it renders them particularly vulnerable to DNA damaging agents such as chemotherapeutics and radiation therapy (5–7). These routine cancer treatments cause chemotherapy-induced gastrointestinal toxicity (CIGT) and radiation induced gastrointestinal syndrome (GIS). CIGT and GIS together affect >80% of cancer patients. They are debilitating pathologies with limited treatment options (8–10). One potential therapeutic avenue would be the development of drugs that protect the TA cells preferentially damaged by these common cancer therapies. However, despite their critical roles in intestinal homeostasis and disease, factors that maintain survival and proliferation of TA cells remain inadequately defined compared to the extensive study of LGR5+ stem cells.

Novel approaches are needed to identify factors that specifically regulate TA cell function. While numerous studies have defined transcriptional control of intestinal stem cells, post-transcriptional regulation of intestinal epithelial homeostasis is only beginning to be understood. Recent research points to important roles for the RNA modification, N6-methyladenosine (m^6^A), in intestinal crypts (11–15). N6- methyladenosine is the most common covalent modification of RNA, occurring on approximately 25% of mRNA transcripts (16, 17). It acts by recruiting RNA-binding proteins that affect mRNA fate, predominantly stability and translation. (18). Global m^6^A methylation patterns in the epithelium can shift with microbial and nutrient contents of the gut and m^6^A-binding proteins have been implicated in intestinal regeneration and the pathogenesis of inflammatory bowel disease (IBD) (11, 13, 15, 19, 20). These studies suggest critical roles for m^6^A in integrating environmental cues with homeostatic and regenerative processes in the intestinal epithelium.

Despite advances in the study of m^6^A in the gut, the effect of global depletion of m^6^A in the intestinal epithelium remains unclear and contradictory. A highly conserved m^6^A “writer” complex installs m^6^A co- transcriptionally in the nucleus of eukaryotic cells. At the core of this complex are the writer proteins Methyltransferase-like 3 and 14 (METTL3 and METTL14) (18). Although METTL3 is the catalytic subunit, both METTL3 and METTL14 are thought to be essential for the methylating activity of the complex (21, 22), and both METTL proteins are deleted interchangeably to define the role of m^6^A in specific tissue or cell types. Recent studies reported essential functions for METTL14 in the survival of LGR5+ stem cells in the colon, but not the small intestine (23, 24). In contrast, another study found that METTL3 deletion caused defects in LGR5+ stem cells in the small intestine (25). However, in the case of METTL3 deletion, rescue of LGR5+ stem cell survival could not rescue tissue homeostasis. Therefore, the primary defect in METTL3- depleted epithelium remains unclear.

In contrast to previous reports emphasizing dysfunction of LGR5+ stem cells, we found that METTL3 deletion induced profound cell death predominantly in small intestinal transit amplifying cells. Disruption of the TA zone was associated with crypt and villus atrophy and widespread reduction in absorptive enterocytes, ultimately resulting in the death of METTL3-depleted mice. Sequencing of m^6^A- modified RNA *in vivo* and ribosome profiling in METTL3-depleted enteroids revealed decreased translation efficiency of methylated transcripts critical to growth factor signaling, including master growth regulator and proto-oncogene, *Kras*. Additional investigation confirmed a novel link between METTL3 and *Kras* methylation and protein expression. Our data identify METTL3 as an essential regulator of intestinal transit amplifying cell survival via direct support of growth factor signaling and KRAS expression. By identifying epitranscriptomic regulation as an indispensable process within TA cells, we highlight a novel gene regulatory mechanism in this critical but poorly understood cell type.

## Results

### Intestinal epithelial METTL3 deletion results in complete growth failure and mortality

To determine the role of METTL3 in intestinal epithelial development and homeostasis, we paired *Mettl3^flox/flox^* mice with the pan-intestinal-epithelial *Villin-*Cre (*Mettl3^VilCreΔ/Δ^*) or its tamoxifen inducible counterpart, *Villin-CreERT2* (inducible *Mettl3^VilCreERΔ/Δ^*). First, we examined *Mettl3^VilCreΔ/Δ^* mice, which had constitutive Cre activation in the small intestinal and colonic epithelium beginning at embryonic day 12.5 (26). These mice were born with Mendelian distribution (X^2^ n=79 p>0.85) and appeared grossly normal at postnatal day 14, as previously described (27). However, from postnatal day 21 to 28, *Mettl3^VilCreΔ/Δ^* mice lost ∼20% starting body weight while controls gained ∼40% (Figure 1A). Body condition and weight loss in *Mettl3^VilCreΔ/Δ^* mice required euthanasia of 70% of mice between postnatal day 16-29 (Figure 1, B-D). To determine whether this phenotype was development-specific, we next examined inducible *Mettl3^VilCreERΔ/Δ^* mice injected with tamoxifen at 8 weeks of age (Figure 2A). After their final tamoxifen injection, inducible *Mettl3^VilCreERΔ/Δ^*mice exhibited an average daily weight loss of ∼2.5% (Figure 2B). Within 10 days, almost all mice experienced critical (>20%) weight loss requiring euthanasia (Figure 2, B and C). These data demonstrate a requirement of METTL3 for growth and survival during the postnatal period and adulthood.

**Figure 1.**
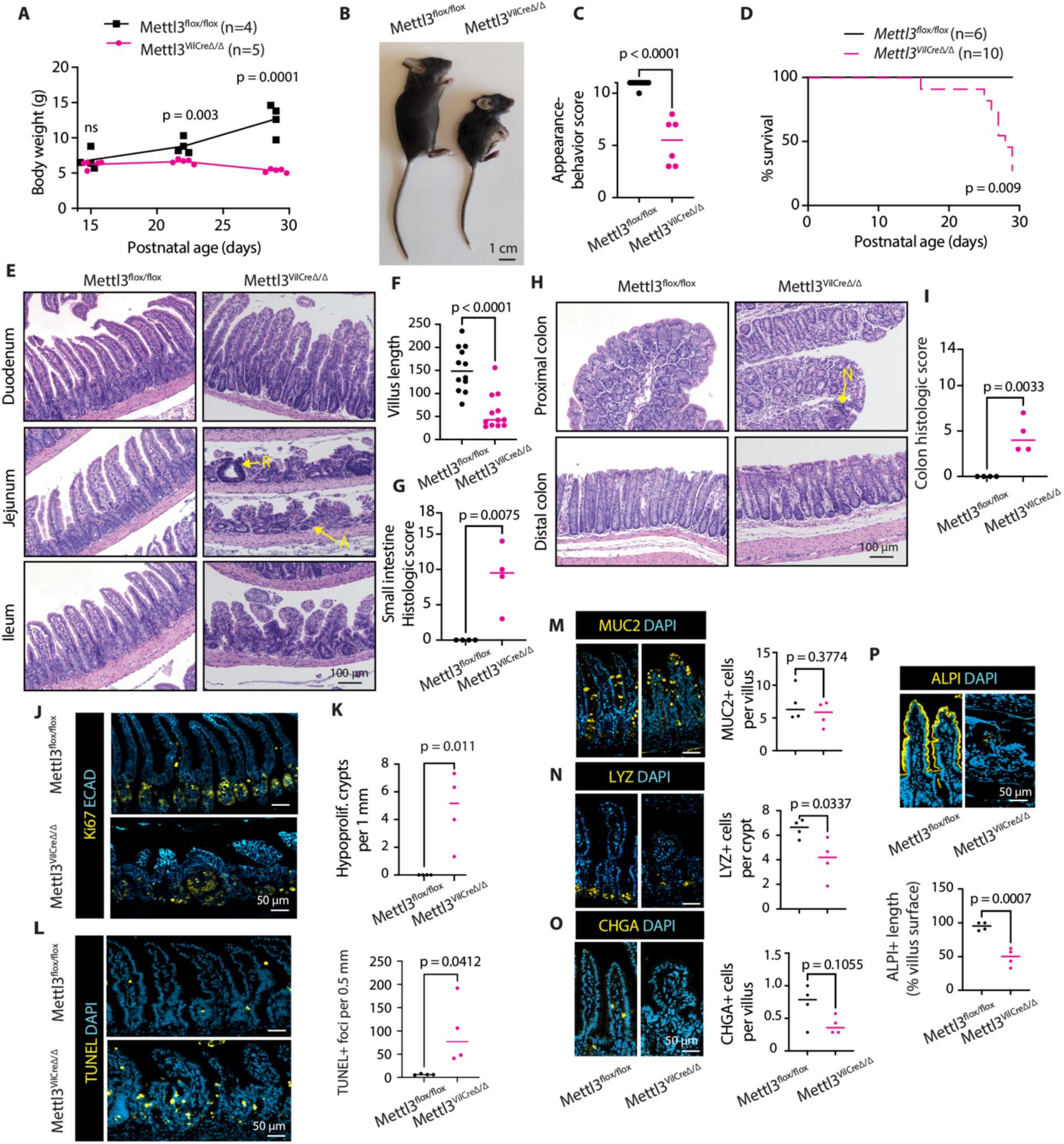
Constitutive METTL3 deletion leads to growth retardation and small intestinal epithelial distortion. **(A)** Growth curves from postnatal day 15 to 29**. (B)** Gross appearance at postnatal day 29. **(C)** Composite appearance and behavior score at postnatal day 29. **(D)** Kaplan-Meier survival curves through postnatal day 29; p value corresponds to Log-rank (Mantel-Cox) test. **(E)** Representative small intestine H&E images. “R” indicates regenerative crypt. “A” indicates atrophic crypt. **(F)** Quantification of jejunal villus lengths, n=4 mice per genotype. **(G)** Composite histological score for small intestine **(H)** Representative colon H&E images. “N” denotes neutrophil invasion. **(I)** Composite histological score for colon **(J)** Representative images of Ki67 in distal small intestine **(K)** Number of hypo-proliferative crypts (<10 Ki67+ cells) crypts per 1 mm distal half small intestine **(L)** Representative images and quantification of TUNEL staining **(M, N, O)** Representative images and quantification of intestinal secretory markers MUC2, LYZ, and CHGA **(P)** Representative images and quantification of alkaline phosphatase. Unless otherwise noted, each data point is the mean of three representative sections imaged per mouse with bar at median value and p value represents unpaired parametric Student’s t test. Immunofluorescence staining and quantification performed in distal half small intestine.

**Figure 2.**
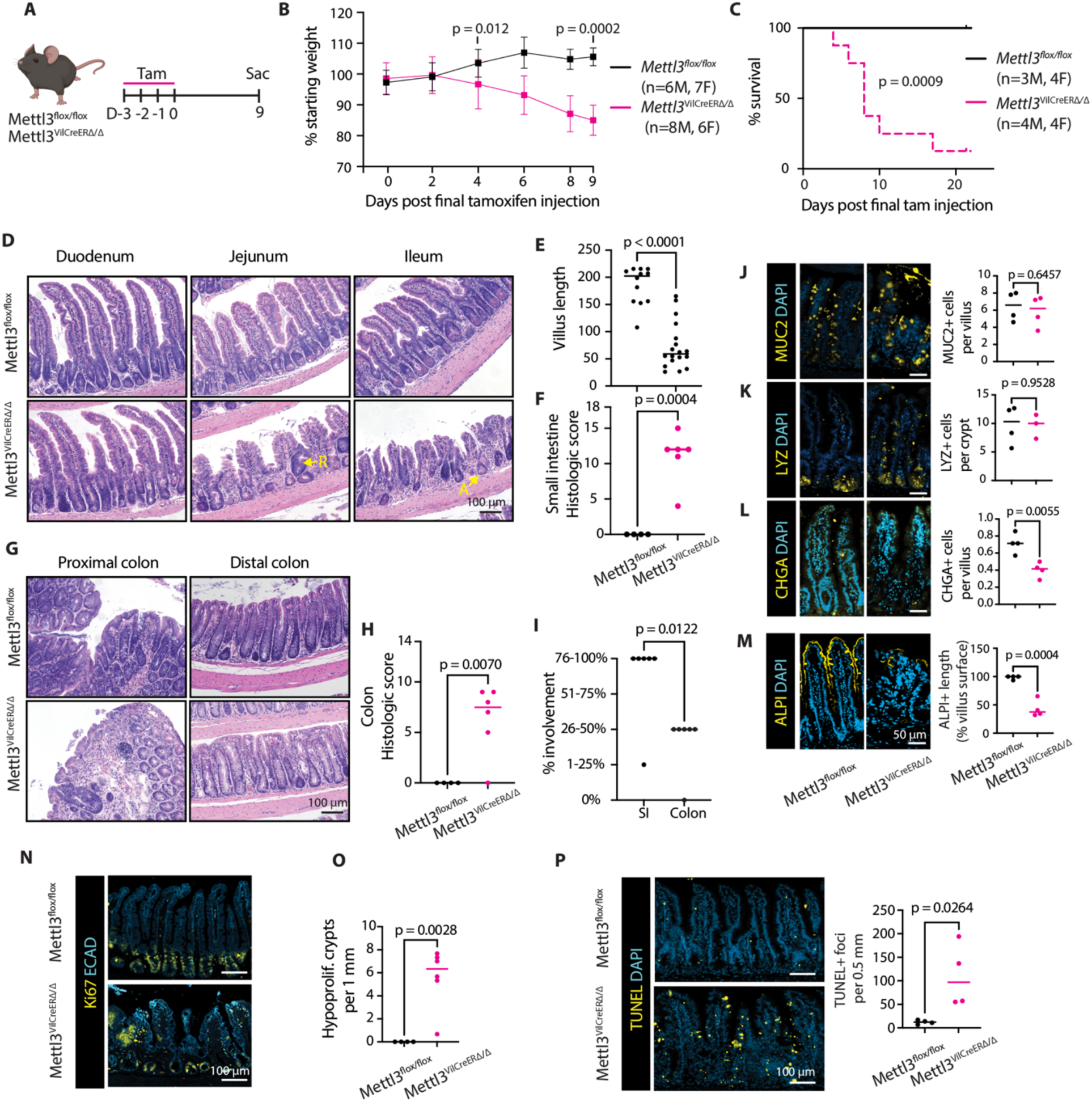
METTL3 is required for survival and small intestinal epithelial homeostasis in adult mice. **(A)** Experimental schematic depicting timing of tamoxifen injection. **(B)** Weight curves through nine days post tamoxifen injection, mean +/- SD. **(C)** Kaplain-Meier survival curves through 22 days post final tamoxifen injecition, p value corresponds to Log-rank (Mantel-Cox) test. **(D)** Representative small intestine H&E images. “R” indicates regenerative crypt. “A” indicates atrophic crypt. **(E)** Quantification of shortest villus lengths in three representative jejunal sections in *Mettl3^flox/flox^* (n=4) and *Mettl3^VilCreERΔ/Δ^* (n=6) mice. **(F)** Composite histological score for small intestine. **(G)** Representative colon H&E images. **(H)** Composite histological score for colon. **(I)** Greatest percent involvement for histological score depicted in F or H. **(J, K, L)** Representative images and quantification of intestinal secretory markers MUC2, LYZ, and CHGA. **(M).** Representative images and quantification of alkaline phosphatase activity. **(N)** Representative images of Ki67 **(O)** Number of hypo-proliferative (<10 Ki67+ cells) crypts per 1mm intestine **(P)** Representative images and quantification of TUNEL staining. Unless otherwise noted, each data point is the mean of three representative sections per mouse with bar at median value and p value represents unpaired parametric Student’s t test. Immunofluorescence staining and quantification performed in distal half small intestine.

### METTL3 deletion induces small intestinal crypt and villus atrophy

We next examined the intestinal pathology of *Mettl3^VilCreΔ/Δ^*and inducible *Mettl3^VilCreERΔ/Δ^* mice to determine the cause of their severe growth failure and mortality. Western blot, *in situ* staining, and m^6^A dot blot confirmed depletion of METTL3 and m^6^A in the intestinal epithelium (Supplemental Figure 1, A-C, and Supplemental Figure 2, A and B). Using a composite histopathological score in both METTL3 knockout models, we noted severe small intestinal defects defined by widespread crypt atrophy alongside villus shortening (Figure 1, E-G, and Figure 2, D-F). We also noted occasional hypertrophic regenerative crypts (Figure 1E and Figure 2D). Although defects were most severe in the distal small intestine, we observed histological changes from the proximal small intestine to the proximal colon. The distal colon was relatively spared (Figure 1, H and I, and Figure 2, G-I). Since METTL3 and METTL14 knockouts are generally considered equivalent, our findings were striking and unexpected given previous studies indicating METTL14 deletion induces severe distal colonic defects but spares the small intestine (23, 24). These data demonstrate that intestinal epithelial METTL3 is required for both the postnatal development and adult maintenance of full-length crypt and villus structures, particularly in the distal small intestine.

### METTL3 is required for intestinal epithelial proliferation and survival

To determine the origins of crypt and villus atrophy, we next evaluated proliferation and apoptosis in *Mettl3^VilCreΔ/Δ^* and inducible *Mettl3^VilCreERΔ/Δ^* mice. Since histological changes were most severe in the distal small intestine, we focused on Ki67 and TUNEL staining in this tissue. In METTL3 knockout epithelium, atrophied crypts previously observed by H&E exhibited drastically reduced Ki67 staining. Where control crypts had an average of ∼30 Ki67+ cells/crypt, METTL3 depleted crypts often exhibited fewer than 10 Ki67+ cells (Figure 1, J and K, and Figure 2, N and O). We also observed some hyperproliferative crypts in both deletion models (>45 Ki67+ cells), although in inducible *Mettl3^VilCreERΔ/Δ^*mice, these were frequently METTL3+ by immunofluorescence (Supplemental Figure 2C). We also identified a >10-fold increase in the mean number of TUNEL positive cells in both *Mettl3^VilCreΔ/Δ^* and inducible *Mettl3^VilCreERΔ/Δ^*mice, demonstrating extensive cell death throughout the villus and crypt (Figure 1L and Figure 2P). These data suggest that both disrupted proliferation and cell survival contributed to epithelial defects in *Mettl3^VilCreΔ/Δ^* and inducible *Mettl3^VilCreERΔ/Δ^*mice.

### METTL3 is required for absorptive enterocyte maturation

To evaluate etiologies of weight loss and epithelial distortion in METTL3 knockout mice, we examined the distribution of differentiated epithelial cells. The differentiated intestinal epithelium comprises secretory and absorptive lineages, which both play essential roles in intestinal homeostasis. Secretory cell depletion can promote epithelial defects due to important roles in producing mucus, antimicrobial compounds, and stem cell niche factors (28, 29). However, even in areas of severe epithelial distortion, we observed maintenance of MUC2+ goblet cells with only minor, inconsistent reductions in LYZ+ Paneth cells and CHGA+ enteroendocrine cells (Figure 1, M-O, and Figure 2, J-L). Alcian blue staining corroborated these findings by indicating mucus production was also maintained in the small intestine, though we did observe reductions in mucus in the proximal colon (Supplemental Figures 1D and 2D). In contrast to modest changes in secretory lineages, we observed a ∼50% decrease in alkaline phosphatase staining in the distal small intestine of both *Mettl3^VilCreΔ/Δ^* mice and inducible *Mettl3^VilCreERΔ/Δ^*mice compared to controls, suggesting dramatic loss of absorptive enterocytes (Figure 1P and Figure 2M). This effect was strongest in areas of severe villus shortening, where we observed almost no alkaline phosphatase staining. Taken together, we find general maintenance of secretory cells but a dramatic reduction in mature absorptive enterocytes in *Mettl3^VilCreΔ/Δ^* mice and inducible *Mettl3^VilCreERΔ/Δ^*mice. We propose that the resulting loss of absorptive capacity may underlie wasting in these knockout mice.

### Alternative METTL3 deletion recapitulates small intestinal defects

We performed the above experiments in inducible *Mettl3^VilCreERΔ/Δ^*mice using *loxP* sites spanning nine exons of the *Mettl3* locus (Supplemental Figure 3A) (30). This large deletion resulted in some reduction of recombination efficiency in the distal colon and to a lesser degree, the small intestine (Supplemental Figure 2, B and C). We were also surprised to identify a small intestinal defect after METTL3 deletion because previously described METTL14 deletion produces no small intestinal phenotype (23, 24). To address these concerns, we tested a second METTL3 deletion model with *LoxP* sites spanning only exon 4 (*Mettl3^VilCreERΔ2/Δ2^*, Supplemental Figure 3B) (31). Inducible *Mettl3^VilCreERΔ2/Δ2^* mice demonstrated efficient deletion of METTL3 in all targeted tissues (Supplemental Figure 3C). In this additional model, we confirmed rapid weight loss and mortality, hypo- and hyper-plastic crypts, preservation of secretory goblet and Paneth cells, and absent distal colonic defects (Supplemental Figure 3, D to G). These data further support the conclusion that METTL3 is essential for small intestinal homeostasis.

### Intestinal defects with METTL3 deletion are independent of intestinal microbiota

Microbial translocation and inflammatory activation are common causes of morbidity and mortality in mice with intestinal epithelial defects. We therefore examined sera of inducible *Mettl3^VilCreERΔ/Δ^*mice for elevated inflammatory cytokines. Intriguingly, there was no significant difference in serum cytokine markers TNF-α, IL-6, IL-1α, IL-1β, IL-10, IL-12 or IFN-γ in *Mettl3^VilCreERΔ/Δ^* mice compared to controls (Figure 3A). Additionally, spleen size—often positively correlated with degree of whole-body inflammation—was reduced (Figure 3B), and small intestine and colon lengths—often reduced in inflammatory conditions— were unchanged (Figure 3, C and D). Taken together, these data suggested a non-inflammatory etiology to morbidity and mortality in *Mettl3^VilCreERΔ/Δ^*mice.

**Figure 3.**
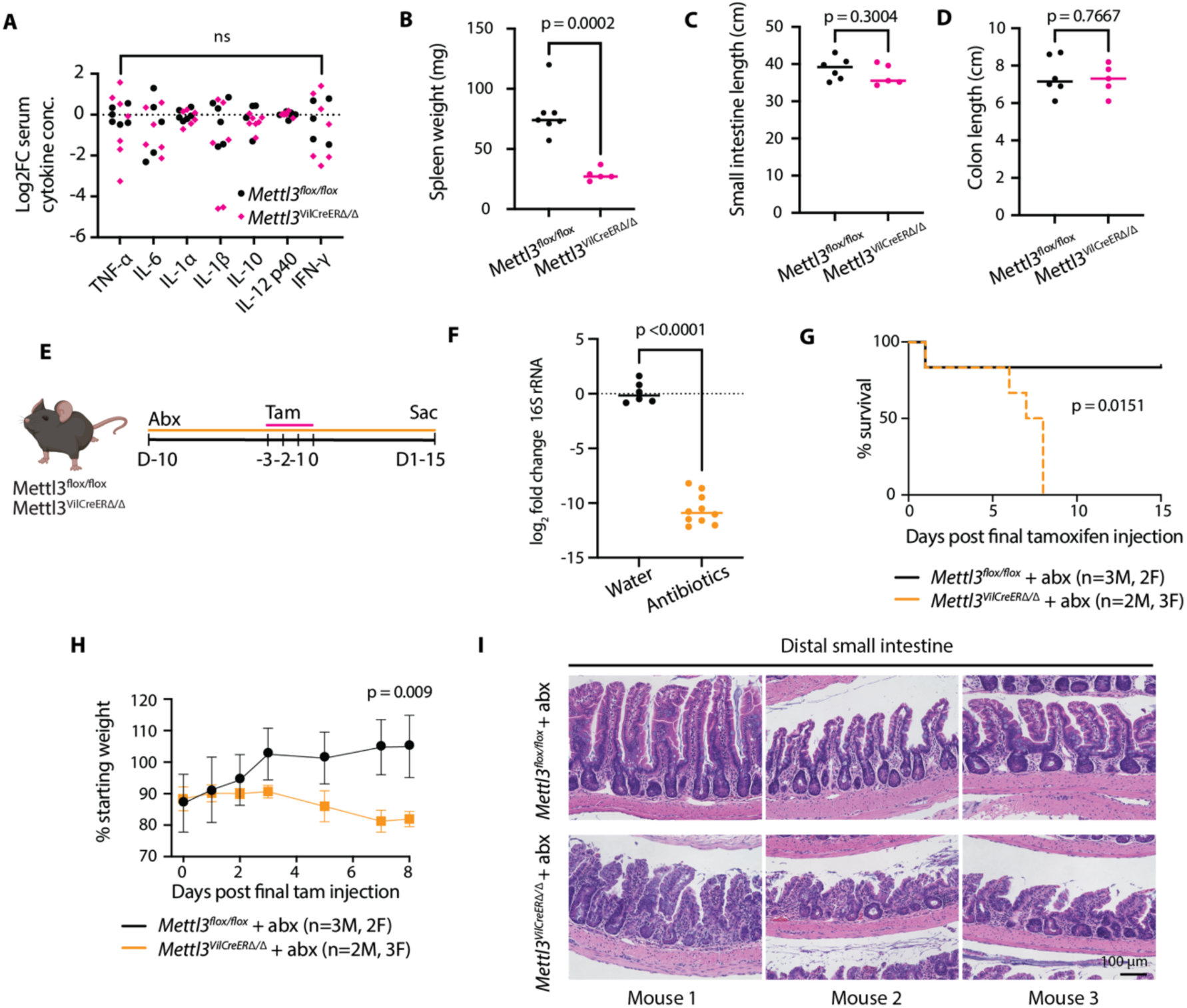
Intestinal distortion and mortality after METTL3 deletion is microbiota independent. **(A)** Log_2_ fold change in serum cytokines in mice nine days post final tamoxifen injection, ns indicates p > 0.05 for all. **(B)** Spleen weight nine days post final tamoxifen injection. **(C, D)** Small intestine and colon length nine days post final tamoxifen injection. **(E)** Experimental schematic for antibiotic (abx) treatment. **(F)** Log_2_ fold change in 16S rRNA amplified from fecal bacterial DNA on final day of tamoxifen injection in antibiotic- treated or water vehicle mice. **(G, H)** Kaplain-Meier survival curve and weight change post final tamoxifen injection in antibiotic treated mice. P-value in (G) represents Log-rank (Mantel-Cox) test. Data in (H) presented as mean +/- SD **(I)** Representative H&E images from matched sections of distal small intestine in n=3 antibiotic treated mice per genotype. Unless otherwise noted, each data point represents a single mouse with bar at median value and p denotes value of unpaired parametric Student’s t test.

Despite no overt inflammatory pathology, the contribution of facility-specific microbiota is a common concern in the evaluation of intestinal phenotypes. We also questioned whether the microbiota contributed to the surprising difference in small intestinal phenotypes between METTL3 and previously described METTL14 knockout intestinal epithelium (23, 24). Therefore, we depleted the microbiota in *Mettl3^VilCreERΔ/Δ^*mice by adding an antibiotic cocktail to their drinking water beginning one week before tamoxifen injection (Figure 3E). Quantitative PCR of genomic 16s rRNA confirmed ∼1000-fold depletion of luminal bacteria (Figure 3F). There was no change in weight loss or mortality in microbiota depleted *Mettl3^VilCreERΔ/Δ^* mice compared to those on normal drinking water (Figure 3, G and H). Histological abnormalities also persisted in microbiota depleted *Mettl3^VilCreERΔ/Δ^* mice (Figure 3I). Taken together, these data suggest minimal contribution of inflammation or microbiota to the gross and histological defects seen in *Mettl3^VilCreERΔ/Δ^* mice. Rather, they further support a model in which loss of absorptive enterocytes diminishes digestive capacity leading to weight loss and mortality in these mice.

### METTL3 deletion immediately triggers transit amplifying cell death

Inducible *Mettl3^VilCreERΔ/Δ^* mice displayed a complex set of histological phenotypes, including adjacent hypertrophic and atrophic crypts and scattered villus and crypt cell death. Some of these changes may be reactive, secondary changes rather than immediately downstream of METTL3 depletion. We therefore examined inducible *Mettl3^VilCreERΔ/Δ^* mice two days after the final tamoxifen injection- the earliest timepoint at which we confirmed METTL3 deletion by immunoblot and *in situ* staining (Figure 4A and Supplemental Figure 2, A and B). At this early deletion timepoint, *Mettl3^VilCreERΔ/Δ^* mice demonstrated a ∼30% increase in crypt height in the small intestine (Figure 4B). This change was associated with increased numbers of Ki67+ cells (Figure 4C) and increased expression of transcripts associated with stem cells and stem progenitors, including key stem cell marker *Lgr5* (Figure 4D). In contrast, we observed profound TA cell death in inducible *Mettl3^VilCreERΔ/Δ^* mice at this same early timepoint. The mean number of TUNEL+ foci per crypt was elevated >20-fold in *Mettl3^VilCreERΔ/Δ^* crypts compared to controls (Figure 4E). Strikingly, almost all TUNEL+ foci in *Mettl3^VilCreERΔ/Δ^* small intestine were in the TA zone, located between the crypt base and the crypt-villus junction. We quantified TUNEL+ staining in the crypt base, TA zone, and villus epithelium and found 3-fold higher cell death rates in the TA zone compared to the crypt base (Figure 4F). These data suggest that the initial defect in inducible *Mettl3^VilCreERΔ/Δ^* mice is widespread transit amplifying cell death.

**Figure 4.**
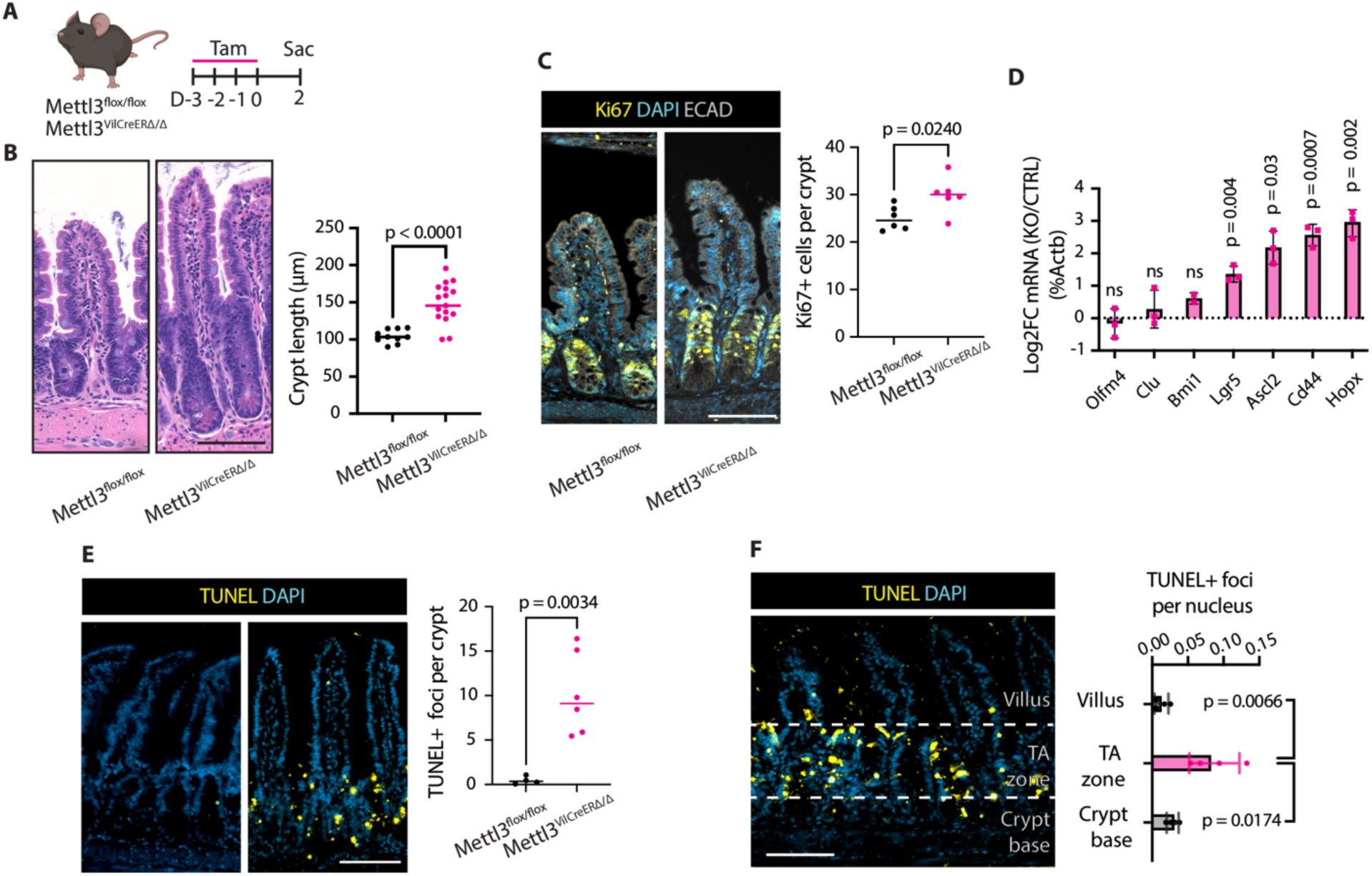
METTL3 deletion rapidly induces death of transit amplifying cells. **(A)** Experimental schematic. **(B)** Representative images and quantification of crypt height. **(C)** Representative images and quantification of Ki67 staining. **(D)** Log_2_ fold change in qPCR quantification of indicated genes normalized to the mean of *Mettl3^flox/flox^* controls and *Actb*. Data from crypt enriched lysates. **(E)** Representative images and quantification of TUNEL staining **(F)** Representative image and quantification of distribution of TUNEL staining in villus, transit amplifying (TA) zone, and crypt base. Unless otherwise noted, each data point is the mean of three representative sections per mouse with bar at median value and p denotes value of unpaired parametric Student’s t test. Scale bar 100 µM. All data from distal small intestine of mice two days post final tamoxifen injection.

### METTL3 deletion causes growth arrest and death in intestinal enteroids

We next wanted to establish an epithelial-specific model for examining the kinetics and mechanism of cell death in *Mettl3^VilCreERΔ/Δ^* mice. We therefore generated 3D organoids from small intestinal crypts (enteroids) and colonic crypts (colonoids) of *Mettl3^VilCreERΔ/Δ^* and *Mettl3^flox/flox^*mice and treated with 4- hydroxytamoxifen (4-OHT) *in vitro* to induce METTL3 deletion (32). We first monitored gross cell death in enteroids and colonoids. We defined dead organoids as those with a complete opaque appearance, leaking of dead cell debris, and absent growth over the subsequent 24 hours. Nearly all *Mettl3^VilCreERΔ/Δ^*enteroids and proximal colonoids died within 5 days post 4-OHT withdrawal (Figure 5, B and C). Only distal colonoids from *Mettl3^VilCreERΔ/Δ^* mice survived 4-OHT treatment, where immunoblotting confirmed deletion of METTL3 (Figure 5D). To map the chronology of enteroid growth and survival after METTL3 deletion, we measured ileal enteroid size and death daily after 4-OHT treatment of *Mettl3^VilCreERΔ/Δ^* and control *Villin-CreERT2* enteroids. Enteroids appeared grossly normal in the first two days after 4-OHT withdrawal. At day 3, *Mettl3^VilCreERΔ/Δ^*enteroids exhibited growth arrest concurrent with increased death, ultimately resulting in complete death by five days post 4-OHT (Figure 5, E-G). In summary, *Mettl3^VilCreERΔ/Δ^* enteroids and colonoids recapitulated tissue-region-specific cell death phenotypes observed *in viv*o. These data confirmed that cell death phenotypes after METTL3 deletion were epithelial-cell-autonomous; they also established enteroids as an effective model of *in vivo* pathologies for further mechanistic investigation.

**Figure 5.**
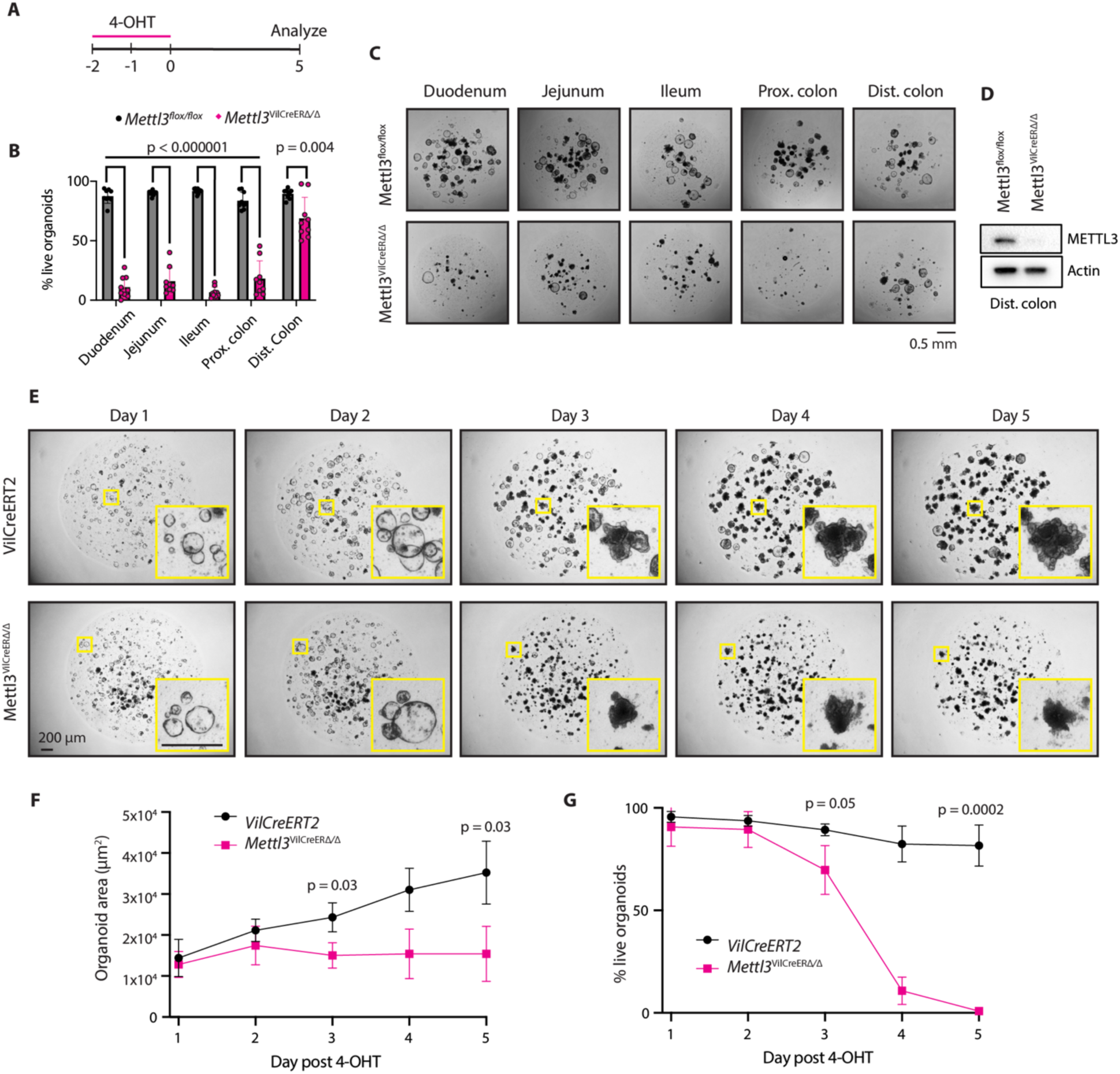
METTL3 deletion triggers growth arrest and death in intestinal epithelial enteroids and colonoids. **(A)** Intestinal epithelial enteroids or colonoids were treated with 1 µm 4-OHT 48 and 24 hours before beginning of time course and analyzed at days 1 through 5. **(B)** Percent live enteroids or colonoids from indicated tissue regions at 5 days post 4-OHT treatment. Each point represents n=9 technical replicates across n=3 passage separated biological replicates per genotype. Bar height at median value with error bars as +/- SD. **(C)** Representative images at 5 days post 4-OHT treatment corresponding to quantification in (B). **(D)** Western blot for METTL3 in surviving *Mettl3^VilCreERΔ/Δ^* distal colonoids six days after 4-OHT treatment. **(E)** Representative images of ileal enteroids in the five days post 4-OHT treatment. **(F, G)** ImageJ quantification of average ileal enteroid 2D area and percent live enteroids in the five days post 4-OHT treatment of *Villin-CreERT2* (VilCreERT2) and *Mettl3^VilCreERΔ/Δ^*enteroids. N=3 passage separated replicates. Data presented as mean +/- SD. P-value represents unpaired parametric Student’s t test at day 3 and day 5.

### Catalytic inactive METTL3 does not rescue death of METTL3 depleted enteroids

Given that METTL3 deletion led to small intestinal epithelial death *in vivo* and *in vitro*, but depletion of the essential methyltransferase co-factor METTL14 has been previously reported to have no effect on small intestinal homeostasis (23, 24), we hypothesized that METTL3 might support intestinal epithelial survival through a non-catalytic mechanism. We reintroduced a non-catalytic METTL3 (DPPW^395-398^ to APPA^395-398^) (33) to *Mettl3^VilCreERΔ/Δ^* ileal enteroids using a lentiviral construct (METTL3^Δcat^). Western and dot blot confirmed rescue of METTL3 expression and depletion of m^6^A in these enteroids (Supplemental Figure 4, A and B). Compared to *Mettl3^VilCreERΔ/Δ^*, *Mettl3^VilCreERΔ/Δ^* + METTL3^Δcat^ enteroids demonstrated delayed cell death through the 4^th^ day post 4-OHT, but still died by day 5 and did not survive to subsequent passages (Supplemental Figure 4, C-E). Previous reports indicated that METTL3 may act independently of METTL14 as a non-catalytic cytoplasmic regulator of protein translation (34–36). However, we observed exclusively nuclear staining of endogenous METTL3 in ileal enteroids, likely precluding this possibility (Supplemental Figure 4F). These data suggest that small intestinal epithelium requires the nuclear, catalytic activity of METTL3 for homeostatic function.

### METTL3 deletion triggers global downregulation of translational efficiency

Previous studies indicate that m^6^A induces global changes in mRNA abundance and translational efficiency (13, 37, 38). To identify METTL3 targets that may mediate phenotypes observed *in vivo* and *in vitro*, we sequenced total RNA and ribosome-protected RNA fragments to assay global changes in RNA abundance and translation after METTL3 deletion. Since METTL3 modifies thousands of transcripts with pleiotropic effects, we elected to analyze *Mettl3^VilCreERΔ/Δ^*ileal enteroids 72 hours after 4-OHT treatment to detect the changes most proximal to METTL3 deletion but prior to widespread cell death. Comparing the approximately 25,000 transcripts detected in both data sets, METTL3 deletion impacted more transcripts at the level of RNA translation than abundance. We found 6,101 transcripts exclusively altered at the level of ribosome protected fragments, 3,004 with changes only in RNA abundance, and 1,374 with changes in both (Figure 6A and Data S1). We next compared the ratio of ribosome protected fragments to mRNA abundance for each transcript to generate the translational efficiency (“TE”) of each transcript (11, 39). Averaging all 2,124 transcripts with significant changes in TE, the predominant effect of METTL3 deletion was reduced TE, with 1,747 transcripts exhibiting reduced TE and 377 with increased TE. Taken together, there was a mean decrease in TE of 39% across all differentially translated transcripts in *Mettl3^VilCreERΔ/Δ^*enteroids compared to controls (Figure 6B). These data suggest a novel role for METTL3 in broadly supporting translation in the small intestinal epithelium.

**Figure 6.**
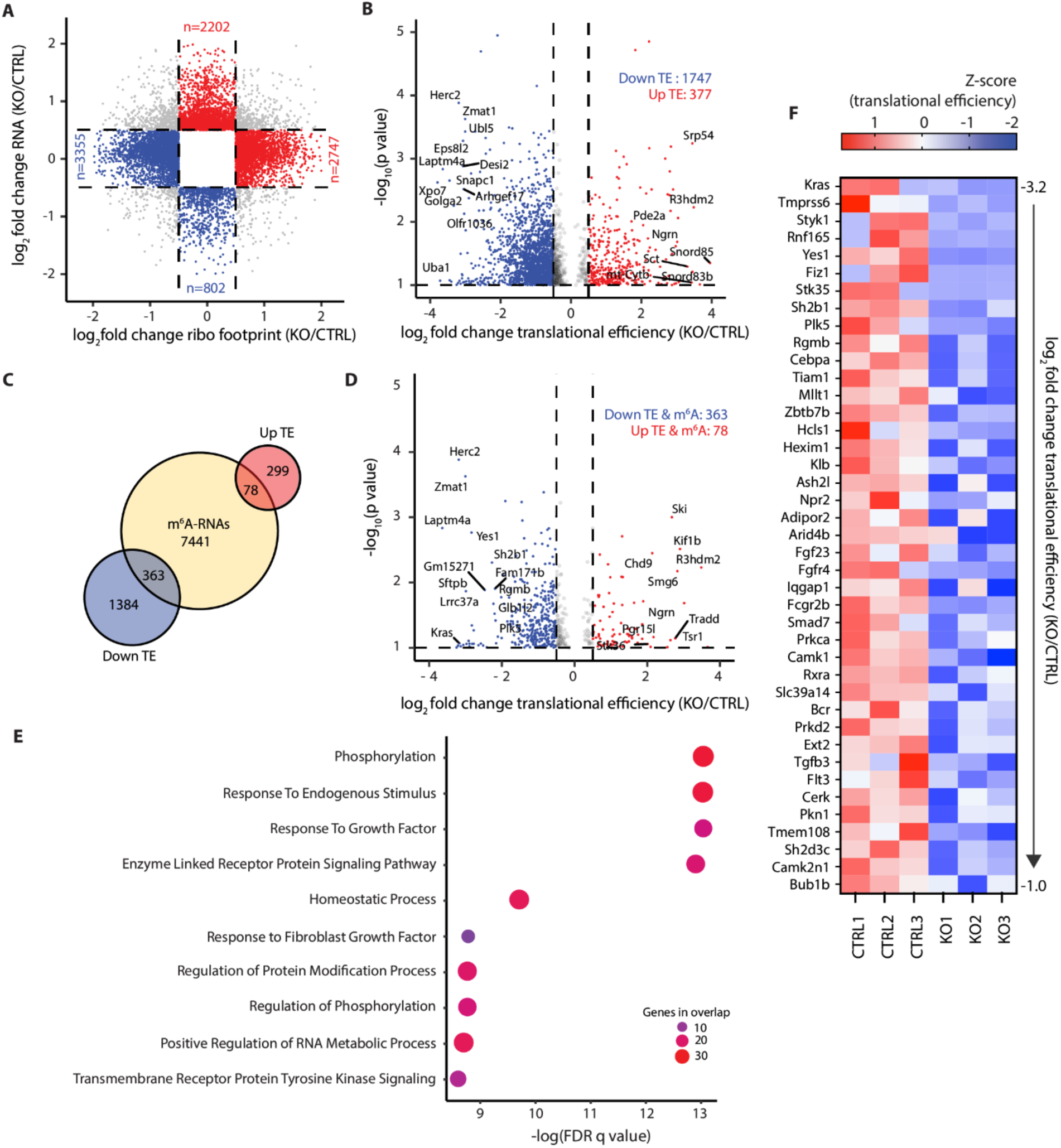
METTL3 deletion leads to a global decrease in mRNA translational efficiency with impacts on growth factor signaling. **(A)** Scatter plot of log_2_ fold change in ribosome footprint and total transcript abundance for all transcripts detected in RNA-seq and Ribo-seq of *Mettl3^flox/flox^* (CTRL) and *Mettl3^VilCreERΔ/Δ^*(KO) ileal enteroids 72 hours post initiation of 4-OHT treatment. Log2 fold change cut off >0.5 or <-0.5. Red marks transcripts increased in RNA or ribosome footprint abundance only. Blue marks decreases in RNA or ribosome footprint abundance only. Grey marks transcripts changed in both. **(B)** Volcano plot of all transcripts with log_2_ fold change in translational efficiency (TE) >0.5 or <-0.5 and -log_10_ p value >1. Red marks all transcripts with increased TE and blue marks all transcripts with decreased TE. **(C)** Venn diagram depicting fraction of transcripts with changes in TE and at least one m^6^A peak as determined by m^6^A-seq of wildtype mouse small intestinal crypt epithelium. Yellow circle is all m^6^A-methylated transcripts detected by m^6^A-seq. Blue circle is all transcripts with decreased TE from (B). Red circle is all transcripts with increased TE from (B). **(D)** Volcano plot of all transcripts displayed in (B), now filtered for transcripts containing at least one m^6^A peak. **(E)** Pathway enrichment analysis comparing transcripts with downregulated TE (log_2_FC < -1) and at least one m^6^A peak against Gene Ontology Biological Process (GOBP) gene sets. Circle color and size both scale with number of genes overlapping between the tested gene set and the GOBP gene set. **(F)** Heat map depicting Z-scores for TE in three *Mettl3^flox/flox^* (CTRL) and three *Mettl3^VilCreERΔ/Δ^* (KO) replicates 72 hours post initiation of 4-OHT treatment. Genes presented are all 42 genes from the four most significantly enriched pathways in (E). Genes are presented in order of greatest decrease in mean TE to smallest. All data from RNA-seq and Ribo-seq of *Mettl3^flox/flox^*(CTRL) and *Mettl3^VilCreERΔ/Δ^* (KO) ileal enteroids 72 hours post initiation of 4-OHT treatment.

### METTL3 deletion downregulates translation of methylated transcripts regulating growth factor signaling

We next performed m^6^A immunoprecipitation and sequencing (m^6^A-seq) in wildtype mouse crypts *in vivo* to define putative direct targets of METTL3 (16, 17, 40). m^6^A-seq yielded 13,763 m^6^A peaks within 7,882 unique transcripts (Data S2). Peaks were distributed across the coding sequence and 3’ untranslated region, with the highest accumulation at the stop codon, consistent with previous reports (Supplemental Figure 5A) (16, 17). We also saw expected patterns of m^6^A enrichment in positive and negative control transcripts (Supplemental Figure 5B). We superimposed m^6^A-seq data onto ribosome profiling data and found that of the 1,747 transcripts with decreased TE after METTL3 deletion, 363 transcripts contained at least one m^6^A peak (Figure 6, C and D). We conducted pathway enrichment analysis on these 363 transcripts using Gene Ontology Biological Process (GOBP) gene sets. We observed the most significant enrichment in transcripts associated with growth factor signaling cascades (Figure 6E). Amongst these transcripts, the largest magnitude reduction in mean TE was for *Kras*, an essential intestinal proto-oncogene that promotes intestinal epithelial proliferation and survival (Figure 6F) (41). Taken together, our ribosome profiling and m^6^A-seq data indicate that METTL3 deletion downregulates translation of methylated transcripts supporting growth factor signaling, including *Kras*.

### METTL3 deletion reduces *Kras* methylation and protein levels and induces cellular senescence

We further explored the putative relationship between METTL3 and KRAS expression by examining m^6^A modification of the *Kras* transcript. Our m^6^A-seq data indicated enriched m^6^A density across the *Kras* gene body, including both UTRs (Figure 7A). To verify m^6^A-seq peaks on the *Kras* transcript and determine their dependence on METTL3 expression, we performed m^6^A-RNA-immunoprecipitation-qPCR (m^6^A-RIP-qPCR) in crypt-enriched lysates from *Mettl3^flox/flox^* and *Mettl3^VilCreERΔ/Δ^* mice three days after final tamoxifen injection. Using qPCR probes targeting both the *Kras* 5’ and 3’ UTR we found significant enrichment of both transcript regions in m^6^A-RIP fractions. However, only m^6^A enrichment in the 3’ UTR appeared dependent on METTL3 expression. The 5’ UTR peak may represent N6,2′-O-dimethyladenosine (m^6^Am), a terminal modification added to the mRNA cap by the methyltransferase PCIF1; m^6^Am is an established off-target of m^6^A antibodies (Figure 7, B and C) (42). These data support the novel finding that METTL3 methylates *Kras* at the 3’UTR.

**Figure 7.**
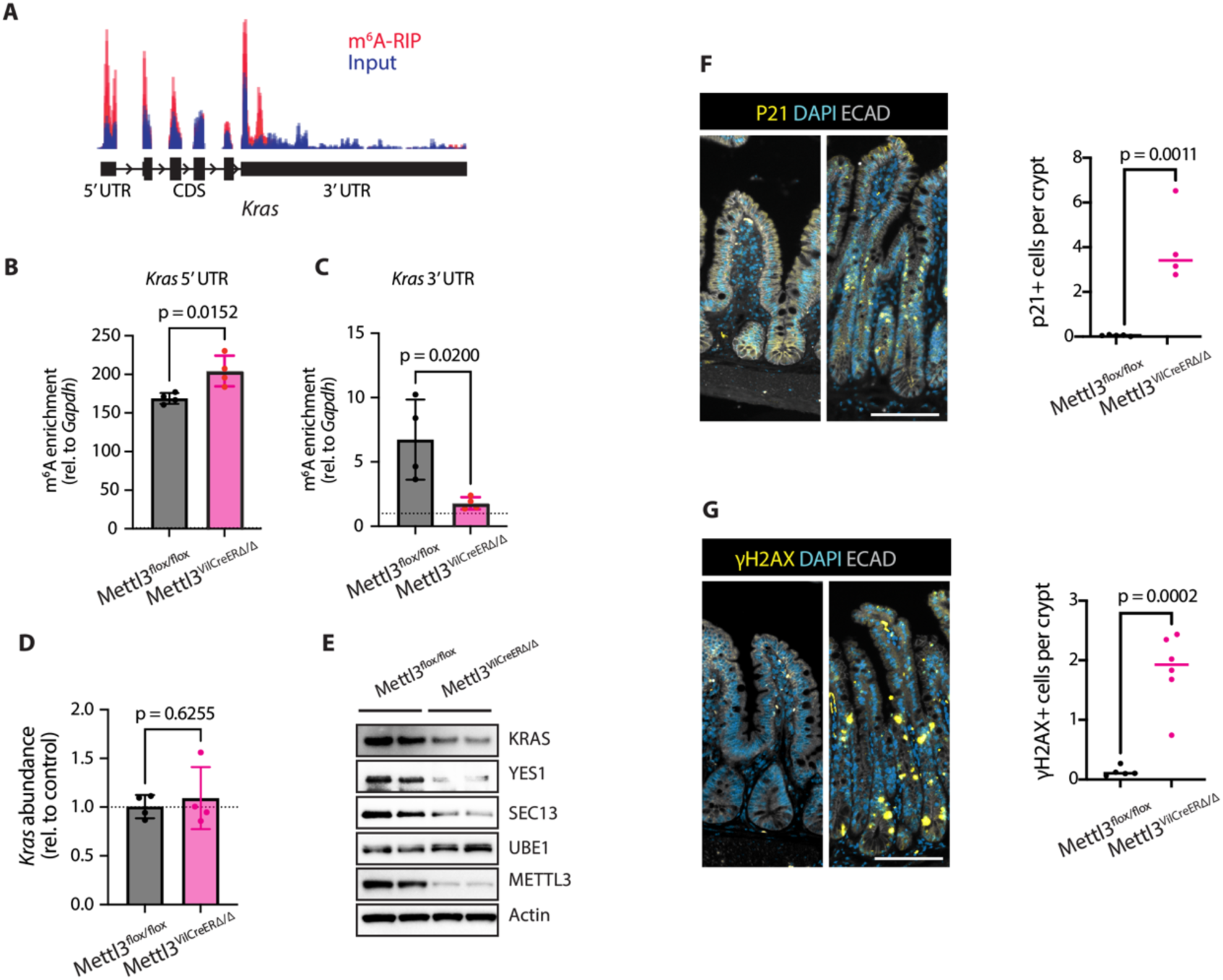
METTL3 deletion downregulates KRAS and induces cellular senescence. **(A)** Integrated Genomics Viewer depiction of read density for m^6^A-RIP (red) and input RNA (blue) for the *Kras* transcript as determined by m^6^A-seq in distal small intestinal crypts of n=3 wildtype mice. **(B, C)** m^6^A-enrichment determined by m^6^A-RIP-qPCR with primers targeting *Kras* 5’ and 3’ UTR in crypt enriched lysates from *Mettl3^flox/flox^* and *Mettl3^VilCreERΔ/Δ^* mice 3 days post-tamoxifen (n=2M, 2F per genotype). The m^6^A enrichment score is the result of dividing the ratio of the target-of-interest in RIP compared to RIP input by the ratio of *Gapdh* (negative control) in RIP compared to RIP input. Data presented as median +/- SD. P-value represents unpaired parametric Student’s t test. **(D)** qPCR for *Kras* transcript in crypt enriched lysates from *Mettl3^flox/flox^* and *Mettl3^VilCreERΔ/Δ^*mice 3 days post-tamoxifen (n= 2M, 2F per genotype). Data normalized to *Actb* and the mean of *Mettl3^flox/flox^* controls. Data presented as median +/- SD. P-value represents unpaired parametric Student’s t test. **(E)** Western blot for top targets with downregulated TE in crypts of *Mettl3^flox/flox^* and *Mettl3^VilCreERΔ/Δ^*mice two days post final tamoxifen injection (n=2 mice per genotype). **(F, G)** Representative images and quantification of p21 and γH2AX staining in distal half small intestine of *Mettl3^flox/flox^* and *Mettl3^VilCreERΔ/Δ^* mice two days post final tamoxifen injection. Images and quantification from areas of most severe histological distortion in distal small intestine of mice two days post final tamoxifen injection. Each data point is the mean of three representative sections imaged per mouse with bar at median value and p denotes value of unpaired parametric Student’s t test. Scale bar 100 µM.

We next examined how methylation of *Kras* impacts its expression *in vivo*. Although METTL3 deletion did not significantly impact *Kras* transcript abundance via qPCR, we observed a decrease in KRAS protein in crypt-enriched lysates from *Mettl3^VilCreERΔ/Δ^* mice compared to controls (Figure 7, D and E). To further validate our Ribo-seq data *in vivo*, we also examined protein expression levels of other top downregulated genes of interest. We confirmed decreases in YES1 (src family kinase and proto-oncogene) and SEC13 (member of the nuclear pore complex implicated in mRNA export) but not UBE1 (primary enzyme in conjugation of ubiquitin) (43–45) (Figure 7E). Taken together, these data suggest that METTL3 promotes KRAS expression without impacting *Kras* transcript levels, providing further mechanistic support for METTL3 as a post-transcriptional regulator of *Kras*.

KRAS is a key mediator of epidermal growth factor (EGF) signaling in epithelial cells (41). Loss of mitogenic signals such as EGF induces cell cycle arrest and senescence in proliferative cells (46). EGF signaling has also been implicated in maintaining genomic integrity in hematopoietic stem cells (47). We therefore assessed senescence and double-stranded DNA breaks in inducible *Mettl3^VilCreERΔ/Δ^*epithelium by staining for p21 and γ-H2AX two days after final tamoxifen injection. *Mettl3^VilCreERΔ/Δ^* crypts demonstrated elevated p21 and γ-H2AX compared to controls, particularly in the TA zone (Figure 7F and 7G). One potential caveat to this finding is the possibility of toxic combined effects of CreERT2 and tamoxifen in intestinal stem cells within a week of tamoxifen exposure (48). To address this, we evaluated p21 and γ- H2AX in tamoxifen-treated *Villin-CreERT2* control mice at the same timepoint. Both p21 and γ-H2AX were significantly increased in *Mettl3^VilCreERΔ/Δ^* mice compared to *Villin-CreERT2* controls (Supplemental Figure 6, A and B). Taken together, these data support the hypothesis that loss of growth factor and KRAS signaling after METTL3 deletion triggers cellular senescence and DNA damage in the crypt TA zone.

## Discussion

Our collective findings examining METTL3 deletion in the intestinal epithelium *in vivo* and *in vitro* support a model in which METTL3 methylates the *Kras* transcript and other transcripts involved in growth factor signal transduction. Methylation promotes translation of these transcripts, maintaining responsiveness to extracellular growth factors and thus maintaining proliferation and survival in the transit amplifying cells of the crypt. In the absence of METTL3, loss of growth factor pathway members such as KRAS leads to cellular senescence and genomic instability. Pathological effects of METTL3 deletion manifest strongest in the TA zone because rapid proliferation demands the expression of proteins involved in transducing growth factor signals. Ultimately, loss of transit amplification reduces crypt and villus size and diminishes absorptive cell maturation without impacting secretory cells (Figure 8).

**Figure 8.**
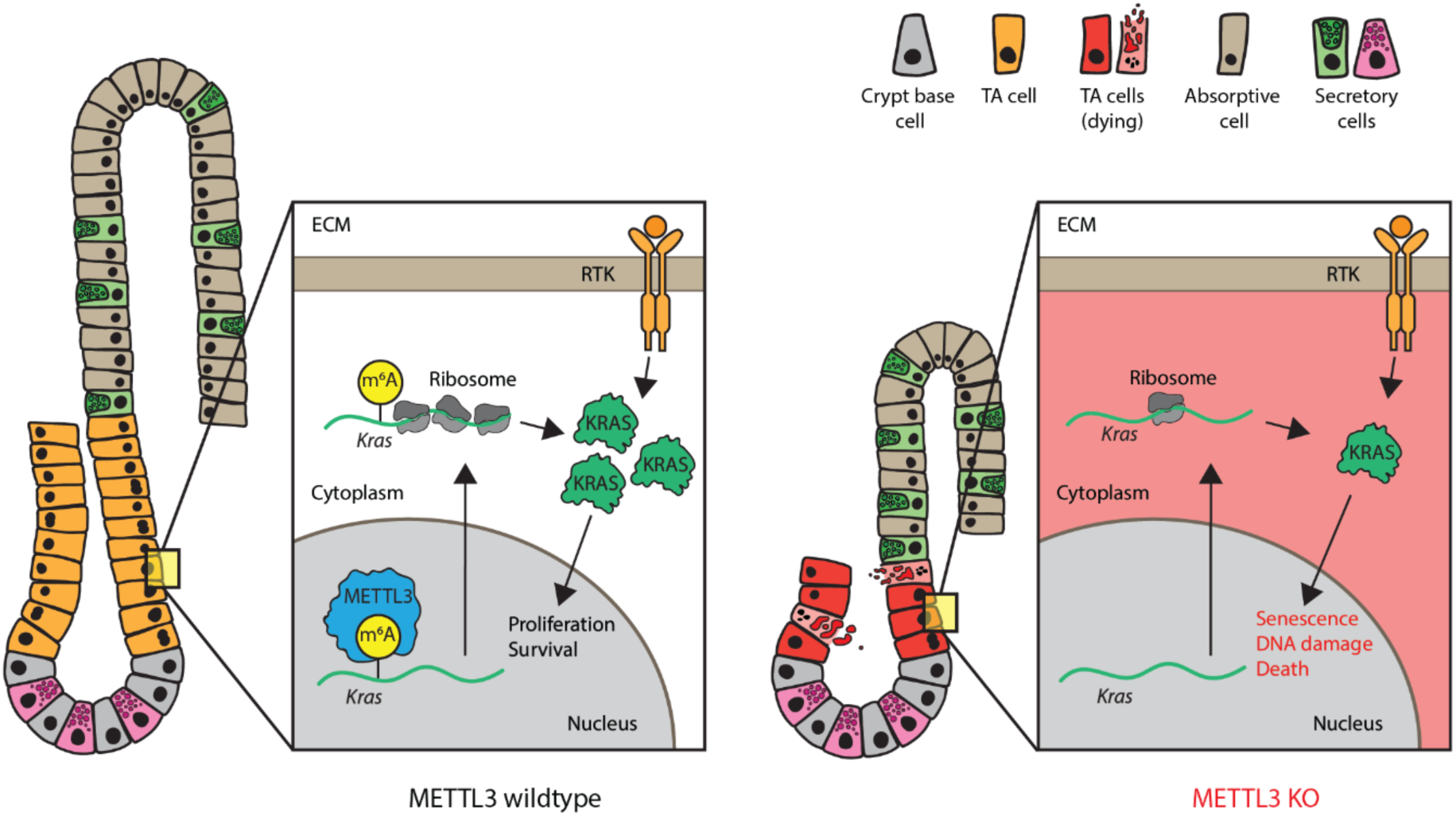
METTL3 maintains growth factor signaling and survival in intestinal transit amplifying cells. Proposed model. METTL3 methylates *Kras* and other transcripts involved in transducing growth factor signaling. Methylation promotes translation of these transcripts, enhancing proliferation and survival in transit amplifying (TA) cells downstream of external growth factors. In the absence of METTL3, a decreased response to extracellular growth factors in METTL3 knockout TA cells leads to cellular senescence and death. Loss of transit amplification results in reduced crypt and villus size and diminished production of absorptive cells. ECM, extra cellular matrix. RTK, receptor tyrosine kinase.

We originally expected that METTL3 deletion would have no overt effect on the small intestinal epithelium because previous reports showed that METTL14 was dispensable in the small intestine (23, 24). While our study does not directly compare METTL3 and METTL14, discrepancies between the present study and previously described METTL14 knockout studies could be due to facility-specific differences (e.g., microbiota). Several factors minimize this possibility. First, two independent groups from the US and China reported colonic but not small intestinal phenotypes with METTL14 knockout, demonstrating persistent phenotypes across mouse facilities. Furthermore, we found small intestinal phenotypes followed METTL3 deletion when testing both constitutive and inducible Cre-drivers, two *Mettl3-floxed* lines, microbiota-depleted mice, and mouse-derived enteroids. The consistency of small intestinal phenotypes spanning numerous orthogonal modes of METTL3 deletion strongly suggests that METTL3 and METTL14 act independently in the small intestine. This unexpected finding suggests novel divergent roles for METTL3 and METTL14 in the gut and underscores the value of examining m^6^A writer proteins in their native context.

Despite their critical role in maintaining homeostatic renewal of the intestinal epithelium, transit amplifying cell dynamics remain poorly understood. We propose that METTL3 is essential for both differentiation and renewal in the transit amplifying zone of the small intestinal crypt. As the site of stem progenitor differentiation, transit amplifying cells are critical to the process of lineage commitment in the cell (2). We observed general maintenance of secretory cells adjacent to areas of atrophic crypts in *Mettl3^VilCre^* and *Mettl3^VilCreERΔ/Δ^* mice. In contrast, these histologically affected areas demonstrated little to no absorptive cell staining. Since mature absorptive cells make up most of the villus surface, transit amplification in the crypt is essential for the production of adequate numbers of absorptive progenitors rather than secretory cells (49). Consistent with our data showing decreased enterocyte maturation concurrent with loss of transit amplifying cells, a recent study suggested a causal relationship between decreased TA proliferation and an increase in the ratio of secretory to absorptive cells (4). Therefore, we propose that METTL3 promotes the production of absorptive cells by maintaining TA cell proliferation.

Inducible METTL3 deletion was also associated with a >20-fold increase in the rate of cell death in the small intestinal crypt, especially in cells of the TA zone. TA cell death immediately followed deletion of METTL3, exceeded rates of cell death in the crypt base 3-fold, and occurred at a timepoint when markers of crypt base stem cells (*e.g., Lgr5*) were preserved or even increased. Therefore, we conclude that the primary, initiating defect in METTL3 depleted epithelium is the death of transit amplifying cells. A recent study posited that TA cells regulate the proliferation of LGR5+ stem cells by controlling the secretion of R-spondins (50). Thus, an initial defect in the TA zone could precipitate the loss of LGR5+ stem cells observed by other groups (25). We also noted increased crypt proliferation at this time of profound proliferative cell death, which initially seemed paradoxical. However, we hypothesize that this increased proliferation represents a regenerative response to increased cell death. Ultimately, the extreme rate of TA cell death appeared to overcome this hyperproliferative response, since most affected crypts exhibited severely diminished proliferation 1-2 weeks after initiating METTL3 deletion. Thus, we conclude that METTL3 specifically regulates the survival of intestinal transit amplifying cells.

Rapid proliferation in the TA zone suggests enhanced dependence on mitogenic factors in these cells. Consistent with this hypothesis, METTL3 deletion downregulated translation of multiple methylated transcripts involved in growth factor signaling, including: growth factor receptor *Fgfr4* (51), src family kinase *Yes1* (43), guanine nucleotide exchange factor *Tiam1* (52), and *Kras*. KRAS is a target of particular interest due to its role as an oncogene in the gut (41), where up to 50% of colorectal cancers harbor a KRAS mutation (53). KRAS binds to guanosine 5’-triphosphate (GTP) in response to extracellular signals such as epidermal growth factor receptor (EGFR) activation. Activated KRAS-GTP then upregulates growth factor targets such as PI3K-Akt, RAF-MEK-ERK, and the GLUT1 glucose transporter (41). Although frequently dysregulated in cancer, these pathways are also essential for homeostatic proliferation, differentiation, and survival (54). Our data support a novel direct relationship between METTL3, *Kras* m^6^A-methylation, and KRAS protein levels. We posit that widescale depletion of growth factor signaling—including but not limited to KRAS—overwhelms cells of the TA zone and leads to cell death with METTL3 deletion.

In summary, we identify METTL3 as an essential post-transcriptional regulator of growth factor signaling and TA cell survival. Our work defines a novel mode of epi-transcriptomic gene regulation in this critical but understudied cell type. This paves the way for future investigation of therapeutic targets that preserve TA cells in the face of DNA damaging agents such as chemotherapeutics and irradiation.

## Materials and Methods

### Animals

*Mettl3^VilCreΔ/Δ^* and *Mettl3^VilCreERΔ/Δ^* were generated by crossing *Vil-Cre 1000* mice (JAX #021504) or *Villin- CreERT2* mice (JAX #020282) to previously described *Mettl3^flox/flox^* mice (30) kindly provided by Dr. Richard Flavell (Yale). *Mettl3^VilCreERΔ2/Δ2^*were generated by crossing *Villin-CreERT2* mice to previously described *Mettl3^flox2/flox2^* mice (31), kindly provided by Dr. Federica Accornero (Ohio State). For the m^6^A- seq experiment, wildtype mice were used (JAX #000664). All mice were C57BL/6J strain and both male and female mice were used. Mice were housed in a temperature-controlled room with 12-hour light and dark cycles and continuous access to food and water.

### Tamoxifen injection, euthanasia criteria, and survival curves

Mice aged 8-9 weeks were injected four times with 50 mg/kg tamoxifen at 10 mg/mL in corn oil at 24-hour intervals. Mice were euthanized once they had reached humane endpoints defined by the IACUC protocol, including: >20% body weight loss, hunched posture, emaciated body condition, or non-responsiveness to stimuli. Survival curves were determined by the number of mice reaching a humane endpoint on each day after tamoxifen injection.

### Microbial depletion

*Mettl3^flox/flox^* and *Mettl3^VilCreERΔ/Δ^* mice were moved to new cages at 7 weeks of age and given sterile DI water supplemented with 0.5g/L Ampicillin (Sigma A9518), 0.5g/L Neomycin (MP Biomedicals 180610), 0.5g/L Gentamicin (Sigma G1914-5G), 0.25g/L Vancomycin (VWR 0990), 0.25g/L Metronidazole (Thermo 210340050), and 4g/L Splenda to enhance taste. After seven days, mice were injected with tamoxifen for four days as described above and stool collected on the final day of injection. Stool from *Mettl3^flox/flox^* and *Mettl3^VilCreERΔ/Δ^* mice on normal drinking water was used as controls. DNA was extracted from stool using the QIAamp Fast DNA Stool Mini Kit (Qiagen 51604) and quantitative PCR was performed as described in the qPCR methods section using primer Ba04230899_s1. qPCR quantification was normalized to stool weight.

### Serum cytokine quantification

Serum supernatant was isolated from the inferior vena cava of euthanized mice and cytokines measured using the Cytometric Bead Array (BD Biosciences) with the amount of capture beads, detection reagents, and sample volumes scaled down tenfold from manufacturer’s protocol. Data was collected on an LSRFortessa flow cytometer (BD Biosciences) with FACSDiva v9.0 (BD Biosciences) and analyzed with FlowJo v10 (BD Biosciences). Statistical outliers were removed in GraphPad Prism v9.3 using ROUT method (Q=1%). Cytokine used were mouse TNFα (BD 558299), mouse IL-6 (BD 558301), mouse IL-1α (BD 560157), mouse IL-1β (BD 560232), mouse IL-10 (BD 558300), mouse IL-12/IL-23p40 (BD 560151), and mouse IFN-γ (BD 558296) with Mouse/Rat Soluble Protein Master Buffer Kit (BD 558266).

### Histology and immunofluorescent staining

Intestines were Swiss-rolled and fixed overnight in 4% paraformaldehyde at 4^°^C prior to processing and embedding. Sections were blocked with Blocking Buffer (1% BSA and 10% donkey serum in PBS) for 1 hour, 25^°^C, before staining with primary antibodies (Supplemental Table 1) at 1:200 overnight at 4^°^C followed by washing with PBS and staining with secondary antibodies at 25^°^C for 25 minutes. Finally, nuclei were counterstained with 1:10,000 DAPI and coverslips mounted with Prolong® Gold Antifade Reagent. Duodenum, jejunum, and ileum were defined as the proximal, middle, and distal third of the small intestine. Proximal colon was defined as the proximal 4cm of the large intestine, and distal colon as the distal 4cm. For alkaline phosphatase staining, we used the Vector® Red Substrate Kit, Alkaline Phosphatase (Vector Laboratories SK-5100) on deparaffinized slides as described in the manufacturer’s protocol. TUNEL was detected using the Click-iT™ Plus TUNEL Assay for In Situ Apoptosis Detection, Alexa Fluor™ 594 dye (Thermo Scientific C10618). All stains were imaged on the Keyence BZ-X100.

### Histopathology and appearance scoring

Histopathological scoring was performed by blinded anatomical pathologist Dr. Benjamin Wilkins based on previously published criteria with an additional score for villus damage for small intestine (Supplemental Table 2) (55). For each category, the recorded score reflects the most severe finding. For percent involvement, the recorded score reflects the length of bowel that was involved by any of the scored processes. Scores were summed for each mouse to generate a separate composite score for small intestine and colon. Mouse appearance/behavior was scored according to a previously described rubric (56).

### Intestinal epithelial organoid cultures

Intestinal sections were splayed open, rinsed in PBS, and rotated at 4°C for 45 minutes in Hank’s Buffered Saline Solution with 10 mM EDTA and 1 mM N-Acetyl-L-cysteine (Sigma A9165). Crypts were isolated by scraping with a glass coverslip followed by vortexing and filtering through a 70 uM (small intestine) or 100 uM (colon) cell strainer. Crypt-enriched suspensions were centrifuged at 500xG, 25^°^C for 5 minutes, washed in PBS, and pelleted again at 500xG, 25^°^C for 5 minutes. Crypts were plated in Matrigel droplets (Corning 354234) and unless stated otherwise, overlaid with the following “ENR” medium: advanced DMEM/F12 media (Thermo 12634028) containing 1X GlutaMAX (Thermo 35050061), 10 mM HEPES (Thermo 15-630-080), 1X Antibiotic Antimycotic (Thermo 15240062), 1X N-2 Supplement (Thermo 17502048), 1X B-27 Supplement (Thermo 17504044), 5 μM CHIR99021 (Cayman 13122), 1 mM N-Acetyl-L-cysteine (Sigma A9165), 50 ng/mL mEGF (PeproTech 315-09), 5% Noggin/R-spondin conditioned medium (generated using protocol from (32)), and 10 µM Y-27632 (LC Labs Y-5301). Colonic epithelial cultures were fed 50% WRN CM media (57). Passage-matched enteroids/colonoids were used for all experiments. For induction of CreERT2, enteroids/colonoids were given 1 µM 4-OHT in 100% ethanol at 48 and 24 hours before the start of the time course then mechanically passaged at day 0.

### Whole mount staining in enteroids

Enteroids were dissociated to single cells using Accutase (STEMCELL Technologies 07920), resuspended in ENR supplemented with 10 µM Y-27632, and plated on chamber slides pretreated with 10% Matrigel in Advanced DMEM/F12. The next day, slides were incubated in 4% PFA at room temperature for 20 minutes and blocked in 10% goat serum in PBS for 30 minutes at 25°C. 1:200 rabbit monoclonal anti-METTL3 (abcam ab195352) and 1:200 goat polyclonal anti-E-Cadherin (R&D Systems AF748) were used at 37°C for 30 minutes, followed by 1:600 Alexa Fluor 488 AffiniPure Bovine Anti-goat IgG (Jackson 805-545-180) and Cy3 AffiniPure Donkey Anti-Rabbit IgG (Jackson 711-165-152) in PBS supplemented with 1:5000 DAPI. Chambers were detached from the chamber slide and coverslips mounted with Prolong® Gold Antifade Reagent.

### qRT-PCR

RNA was isolated using the Quick-RNA Miniprep Kit (Zymo R1054). RNA was reverse transcribed using MultiScribe™ Reverse Transcriptase (Thermo Scientific 4311235) with random hexamer primers.

Quantitative PCR was performed using the Applied Biosystems TaqMan Fast Advanced Master Mix (Thermo Scientific 4444556) with TaqMan™ Gene Expression Assay (FAM) primers (Supplemental Table 3) on an Applied Biosystems QuantStudio 3.

### Western blotting

Cells were lysed using RIPA buffer (CST 9806S) supplemented with 1:100 protease phosphatase inhibitor (CST 5872S) and protein concentration was measured using the Pierce™ BCA Protein Assay Kit (Thermo 23225). Lysates were boiled for 5 minutes with 100 mM DTT and 1X LDS buffer (GenScript M00676) and run on NuPAGE 4-12% Bis-Tris gels (Thermo NP0335BOX) in 1X MOPS-SDS buffer (BioWorld 10530007-2) or 1X MES-SDS buffer (Thermo NP0002, use for low MW proteins). Gels were transferred onto PVDF transfer membrane in 2X NuPAGE transfer buffer (Thermo NP006) and membranes blocked for 1 hr in 5% milk in TBS-T (TBS, 0.1% Tween-20), and then placed in 1:1000 primary antibody (Supplementary Table 1) overnight in 5% milk in TBS-T at 4^°^C. Blots were washed in TBS-T and placed in 1:2000 secondary antibody (Supplementary Table 1) for 1-2 hours at 25^°^C. Blots were then subjected to Supersignal® West Femto Maximum Sensitivity Chemiluminescent Substrate and imaged on the Biorad Gel Doc XR.

### m^6^A dot blot

mRNA was isolated using the Dynabeads™ mRNA DIRECT™ Purification Kit (Thermo 61011). >30 ng of mRNA was heated to 65°C for 2 minutes, placed on ice for >5 minutes, and then pipetted onto a Hybond N+ nitrocellulose membrane and exposed to 254 nm UV light for 5 minutes. The nitrocellulose membrane was then washed in TBS-T (TBS, 0.1% Tween-20). Blots were then incubated in primary and secondary antibody and developed as described in the section on Western blotting. We detected m^6^A using anti-m^6^A primary antibody CST 56593. Total blotted mRNA was stained using 0.04% Methylene Blue (LabChem LC168508) in 0.5M sodium acetate.

### RNA-seq and Ribo-seq

Ileal enteroids were expanded in Matrigel in 50% WRN media until 8 to 10 million cells per replicate was achieved. Passage separated replicates from the same enteroid line were used. Media was supplemented with 2 uM 4-OHT 72 hours prior to collection. Enteroids were collected by mechanical dissociation, pelleted, and snap frozen in liquid nitrogen. Frozen enteroid pellets were resuspended in ice-cold lysis buffer containing 20mM Tris-HCl pH 7.4, 150mM MgCl2, 150mM NaCl, 100ug/ml cycloheximide, 1% v/v Triton X-100, 1mM DTT, 1U/ul SUPERase·IN RNase inhibitor (ThermoFisher), 25U/ml Turbo DNase1 (ThermoFisher), and 1X EDTA-free protease inhibitor cocktail (Roche). Cells were lysed by trituration through a 26-gauge needle 10 times. Samples were processed and libraries prepared as previously described (39) with the following modifications. First, we performed sucrose cushion to pellet the ribosome-associated mRNAs and proceeded with RNAse I digestion (10 U/μl by the Epicentre definition) for 30ug of RNA. Samples were incubated at room temperature for 45 minutes with gentle agitation, and the digestion was quenched by adding 200 units of SUPERase·IN. Then, we directly extracted the ribosome-protected fragments (RPF) using the TRIzol reagent and performed gel size selection of the RPFs of 15-35 nucleotides in length. Second, we performed rRNA depletion using RiboCop for Human/Mouse/Rat kit (Lexogen). Once the rRNA-depleted RPF fragments were obtained, they were pooled together, and the libraries prepared with Unique molecular identifiers (UMIs) for deduplication. The multiplexed library was then sequenced on Illumina HiSeq 4000 with PE150 runs (paired end reading of 150 bases), with a sequencing depth of 60 million raw reads/sample. For each sample, one-tenth (150ul) of the lysate was saved for RNA-seq. For RNA-seq, total RNA was extracted using TRIzol LS reagent (Ambion) and purified using Quick-RNA Microprep kit (Zymo) following the manufacturer’s protocol. Libraries were prepared from total RNA using the Smarter® Stranded Total RNA-Seq Kit v2 - Pico Input Mammalian (Takara Bio 634411) and sequenced on a Novaseq 6000, SP Reagent Kit 1.5 (100 cycles). Raw sequencing data were demultiplexed, adaptors were removed using cutadapt (58), contaminant sequences (rRNA and tRNA) were depleted, and reads were deduplicated (umi_tools dedup). Reads were aligned to all transcripts on the mouse reference chromosomes (Gencode version M31, GRCm39) using Kallisto (59). Translation efficiency (TE) was calculated by dividing the TPM in the total RNA library by the TPM in the RPF library for each individual transcript and sample. Pathway enrichment analysis was performed using mouse gene symbols orthology-mapped to the human genome and tested against Gene Ontology Biological Process (GOBP) gene sets with the Molecular Signatures Database (gsea-msigdb.org).

### m^6^A-seq

Three wildtype C57BL/6J mice (Jax #000664, 2 male 1 female) aged 8 weeks were used. Crypts were isolated from the distal half of the small intestine as described above and dissociated in 10% FBS in 1X PBS supplemented with 20 μg/mL Liberase TH (Roche 05401135001), and 35 μg/ml DNaseI (Roche 10104159001) for 20 minutes at 37^°^C with frequent agitation. Cell suspensions were then stained with 1:200 PE-EpCAM (BioLegend 118205), 1:200 FITC-CD45 (BioLegend 103107), and 1:1000 DRAQ7 (Thermo Scientific D15106) on ice for 30 minutes in the dark. ∼800K live epithelial cells per mouse were then isolated by flow cytometry by sorting for DRAQ7- CD45- PE+ cells directly into Trizol LS (Ambion 10-296-010) on a FACSJazz Sorter. RNA was isolated from Trizol LS using the Direct-zol RNA Microprep Kit (Zymo R2062). RIP-seq was performed according to the “Refined RIP-seq” protocol for low input material (40) with the following specifications: 2 µg anti-m^6^A antibody (Synaptic Systems 202 003) was conjugated to magnetic beads and incubated with 6 ug total RNA that was previously fragmented to ∼300 nt fragments with 10 mM ZnCl2 in 10 mM Tris-HCl for 4 minutes at 70°C (5% of fragmented RNA was set aside as input). After 2 hours immunoprecipitation at 4°C, RNA-bead complexes were washed in high and low salt buffer and RNA was eluted from the RNA-bead complexes using the Qiagen RNeasy Plus Micro Kit (Qiagen 74034). Isolated m^6^A-enriched RNA was immunoprecipitated a second time using 2 µg of a second anti-m^6^A antibody (Abcam ab151230). Final cDNA Libraries were prepared from twice immunoprecipitated RNA and fragmented input total RNA using the Smarter® Stranded Total RNA-Seq Kit v2 - Pico Input Mammalian (Takara Bio 634411) and sequenced on a Novaseq 6000, SP Reagent Kit 1.5 (100 cycles). Raw reads were aligned to mm10 (gencode_M23_GRCm38.p6) using STAR aligner (2.7.9a). N6-methyladenosine peaks were identified using exomePeak2 (version 1.2.0), followed by a metagene analysis using the bioconducter packages Guitar_2.10.0 and TxDb.Mmusculus.UCSC.mm10.knownGene_3.10.0.

### m^6^A-RIP-qPCR

For m^6^A-RIP-qPCR, four *Mettl3^flox/flox^* and four *Mettl3^VilCreERΔ/Δ^*mice aged 8 weeks were used (2 male, 2 female mice per genotype). Intestinal crypts were isolated from the distal half of the small intestine as described above but without digestion to single cells. Whole crypts were pelleted, resuspended in TRI Reagent (Sigma 93289), and RNA was isolated using the Direct-zol RNA Miniprep Kit (Zymo R2050). Next, 20 µg of total RNA was fragmented and immunoprecipitated twice according to the “Refined RIP-seq protocol” as described above. Immunoprecipitated RNA and fragmented input total RNA was reverse transcribed using the High-Capacity cDNA Reverse Transcription Kit (Thermo 4368814) and qPCR was performed as described in the “qRT-PCR” section. For calculation of m^6^A enrichment, first we calculated the %input for every target in every RIP sample as 2^-(Ct of target gene in RIP – Ct of target gene in input sample). Then we calculated m^6^A enrichment relative to *Gapdh* by dividing the %input of the target of interest (e.g., *Kras 5’ or 3’UTR)* by the %input of *Gapdh* for each sample.

### Lentiviral constructs and transduction

Accutase-digested (STEMCELL Technologies 07920) enteroids were resuspended in ENR with 1:4 lentivirus solution, TransDux MAX Lentivirus Transduction Enhancer (System Biosciences LV860A-1), and 10 uM Y-27632 and plated as monolayers on Collagen I (Advanced Biomatrix 5010). Lentiviral constructs and prepared virus were generated by VectorBuilder. The transfer vector for catalytic inactive METTL3 expression contained a CMV-EGFP:T2A:Puro selection cassette and the mPGK promoter upstream of the *Mus musculus Mettl3* ORF edited at positions 1183-1194 from GACCCACCTTGG to GCCCCACCTGCG yielding DPPW -> APPA synonymous mutation in the catalytic site as previously described (33). The “GFP” control vector contained a CMV-mCherry:T2A:Puro selection cassette and the EGFP ORF under control of the CMV promoter. After overnight incubation with lentivirus, enteroid monolayers were mechanically dissociated and replated as 3D enteroids in Matrigel and selected for antibiotic resistance genes after 48 hours.

### Quantification and statistical analysis

Unless otherwise noted, quantification of immunofluorescent staining was performed using three representative images taken per mouse using a 20X objective. Images were taken in three areas of most severe histological distortion (defined as a high-powered field with greatest changes in crypt and/or villus morphology) in distal half of the small intestine of tested mice. Three representative sections were chosen from matching regions of the distal half of the small intestine in control mice. Quantification was performed by individuals blinded to mouse genotype. Unless otherwise noted, each graphed data point is the mean of three quantified representative images per mouse. P values were calculated with the unpaired parametric Student’s t test in GraphPad Prism. P values for survival curves were calculated using the Log-rank Mantel-Cox test in GraphPad Prism.

### Study approval

Mouse experiments and handling were approved under IACUC protocol 001278 at the Children’s Hospital of Philadelphia.

## Supporting information

BioRender License

## Data and materials availability

All data needed to evaluate the conclusions in the paper are present in the paper and/or the Supplementary Materials. Raw sequencing data are uploaded to Dryad at https://doi.org/10.5061/dryad.5tb2rbp8s.

## Author contributions

Conceptualization: CHD, KEH

Methodology: CHD, EAM, SV, PS

Formal analysis: CHD, KEH^2^, SV, YZ, BJW

Investigation: CHD, KEN, SV, RM, SKN, KK, LP, XM, AC

Visualization: CHD, KEH^2^

Supervision: CHD, KEH, PS, MDW

Writing—original draft: CHD, KEH

Writing—review & editing: CHD, KEH, MDW, KEN, KK

KEH: Katharyn E. Hamilton

KEH^2^: Katharina E. Hayer

## Acknowledgments

We thank Dr. Richard Flavell (Yale University), Dr. Federica Accornero (Ohio State University), and Dr. Brian Capell (University of Pennsylvania Perelman School of Medicine) for generously providing Mettl3- floxed mice used in this study. For feedback, technical assistance, and/or sharing of laboratory reagents, we thank: Dr. Michael Silverman, Dr. Michael Abt, Dr. Tatiana Karakasheva, Gloria Soto, Dr. Amanda Muir, Dr. Masaru Sasaki, Joshua Wang, Dr. Lan Lin, Dr. Samir Adhikari, Dr. Derek Sung, and Nora Kiledjian (Children’s Hospital of Philadelphia/University of Pennsylvania Perelman School of Medicine) and Dr. Igor Brodsky, Dr. Chris Lengner, and Zvi Cramer (University of Pennsylvania School of Veterinary Medicine). We also thank the following scientific cores and centers: The Center for Molecular Studies in Digestive and Liver Diseases (P30DK050306) and the Molecular Pathology and Imaging Core (RRID: SCR_022420), the Flow Cytometry Core at the Children’s Hospital of Philadelphia, and the Center for Applied Genomics at the Children’s Hospital of Philadelphia. Artwork depicting mouse injection schemes was created with BioRender.com

## Supplementary Materials

**Supplemental Figure 1.**
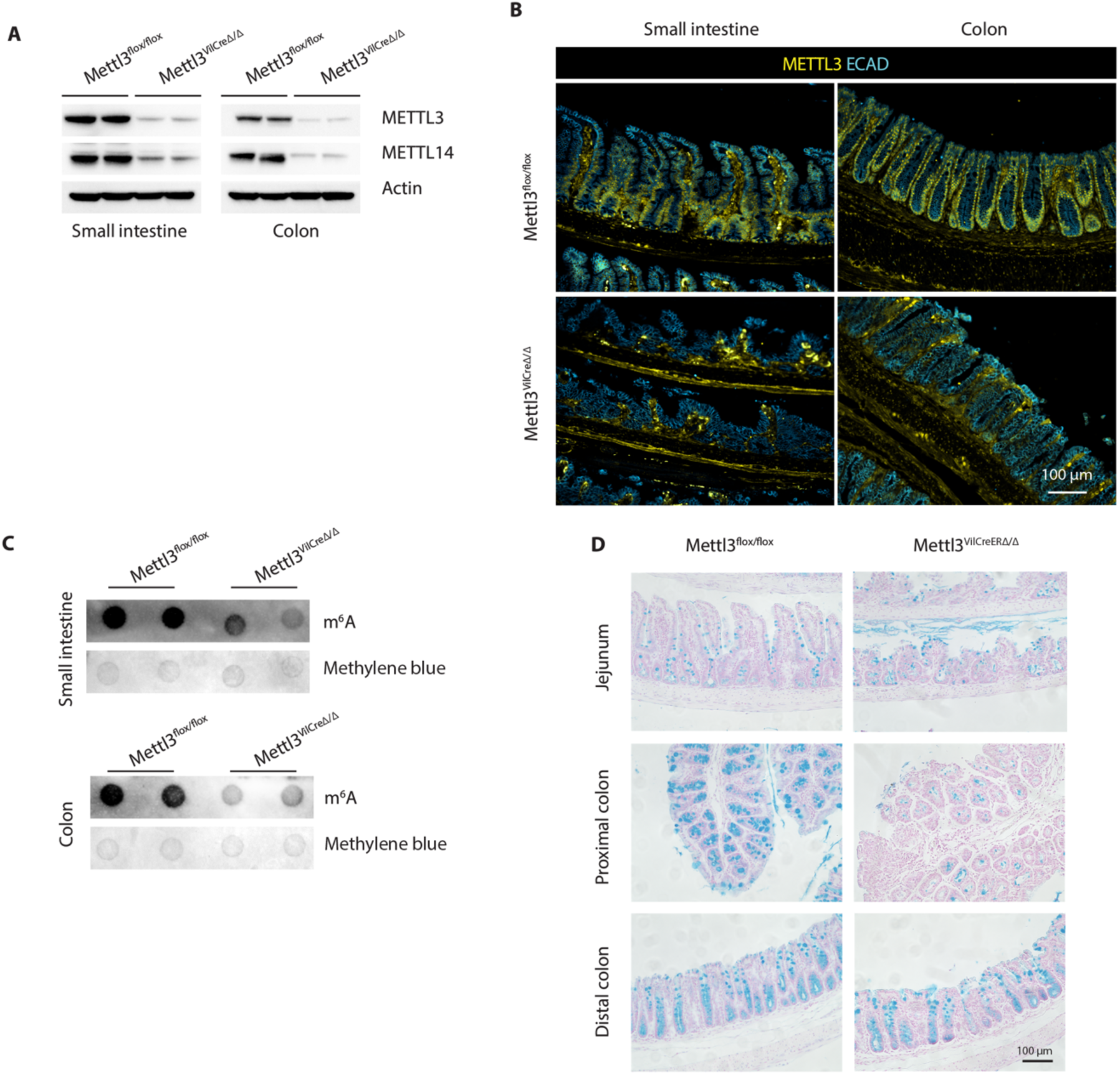
Verification and additional intestinal phenotypes in *Mettl3^VilCreΔ/Δ^* mice. **(A)** Western blot for METTL3 and METTL14 in epithelial crypt enriched lysates from distal half of small intestine and colon. **(B)** Immunofluorescent staining of METTL3 in jejunum. **(C)** m^6^A dot blot in isolated crypts of distal half of small intestine and colon. **(D)** Representative Alcian blue staining in small intestine and colon. All data from postnatal day 29.

**Supplemental Figure 2.**
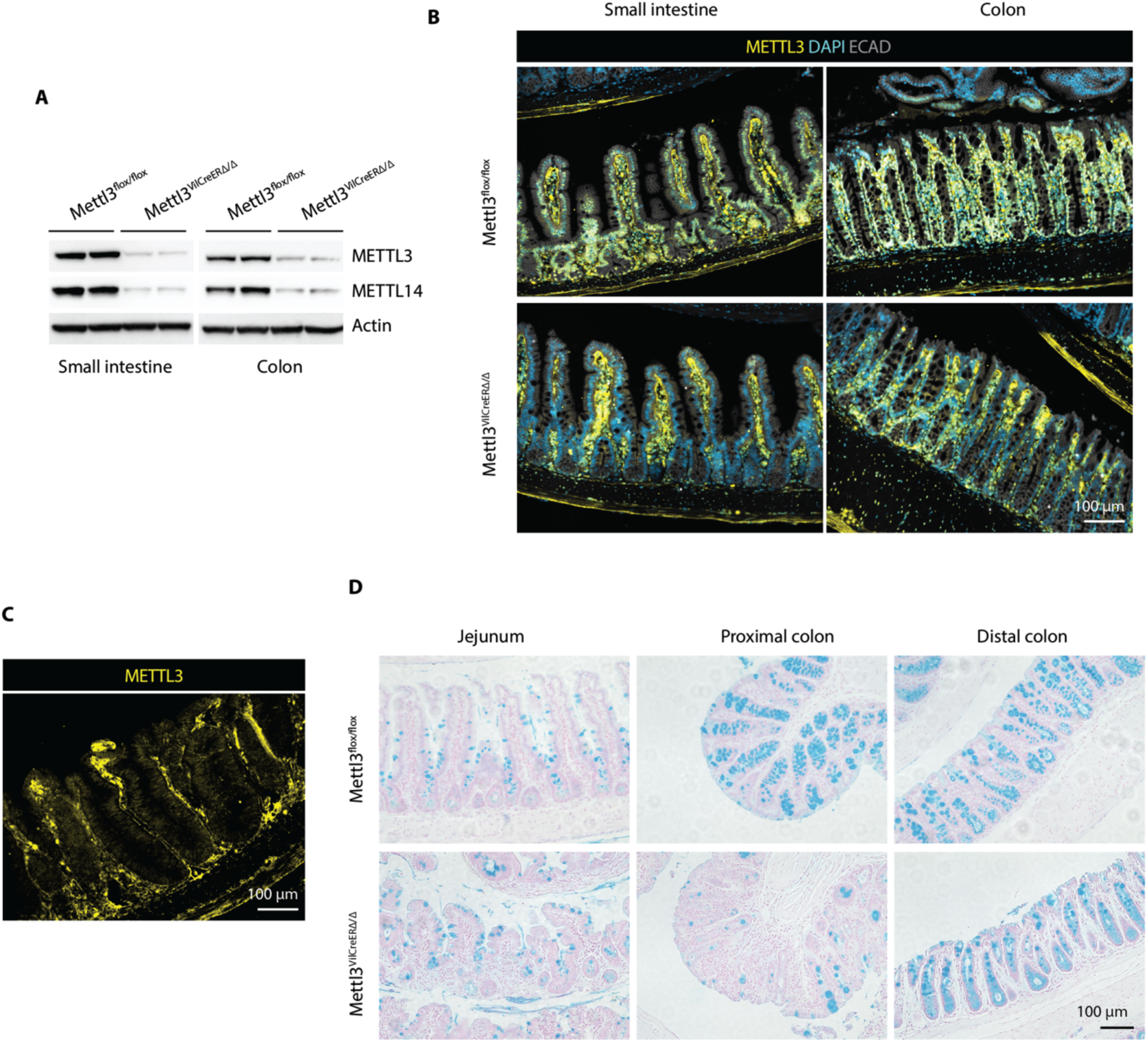
Validation and additional characterization of inducible *Mettl3^VilCreERΔ/Δ^* mice. **(A)** Western blot for METTL3 and METTL14 in epithelial crypt enriched lysates from distal half of small intestine and colon in mice two days post final tamoxifen injection. **(B)** Immunofluorescent staining of METTL3 in jejunum and colon two days post final tamoxifen injection. **(C)** METTL3 staining in hypertrophic small intestinal crypts in a *Mettl3^VilCreERΔ/**Δ**^* mouse nine days post final tamoxifen injection. **(D)** Representative Alcian blue staining nine days post final tamoxifen injection.

**Supplemental Figure 3.**
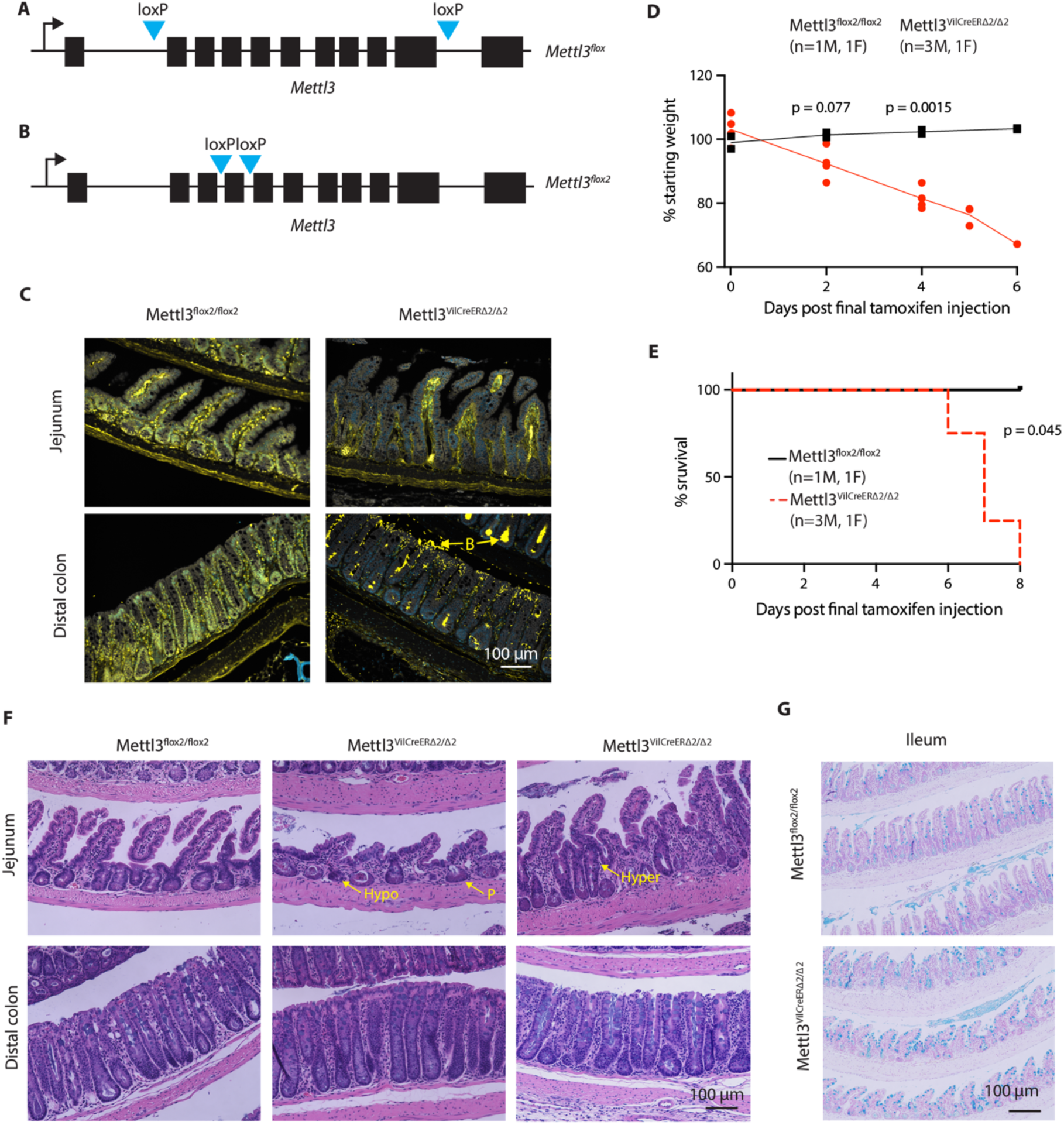
A second *Mettl3* deletion model recapitulates intestinal phenotypes. **(A, B)** Schematic depicting *loxP* sites in the *Mettl3* gene. **(C)** METTL3 staining in jejunum and distal colon. “B” indicates presumed off-target luminal bacterial staining with anti-METTL3 antibody. **(D)** Weight loss post final tamoxifen injection in *Mettl3^flox2/flox2^* (n=2) and *Mettl3^VilCreERΔ2/Δ2^* (n=4) mice. Each individual point represents one mouse. **(E)** Kaplain-Meier survival curve post final tamoxifen injection in *Mettl3^flox2/flox2^* (n=2) and *Mettl3^VilCreERΔ2/Δ2^*(n=4) mice. **(F)** Representative H&E images from small intestine and colon. “Hypo” indicates hypoplastic crypts. “Hyper” indicates hyperplastic crypts. “P” indicates crypts dominated by Paneth cell granules. **(G)** Representative Alcian blue staining of ileum. Images from areas of most severe histological distortion in distal small intestine of mice meeting euthanasia criteria or littermate, tamoxifen-injected controls.

**Supplemental Figure 4.**
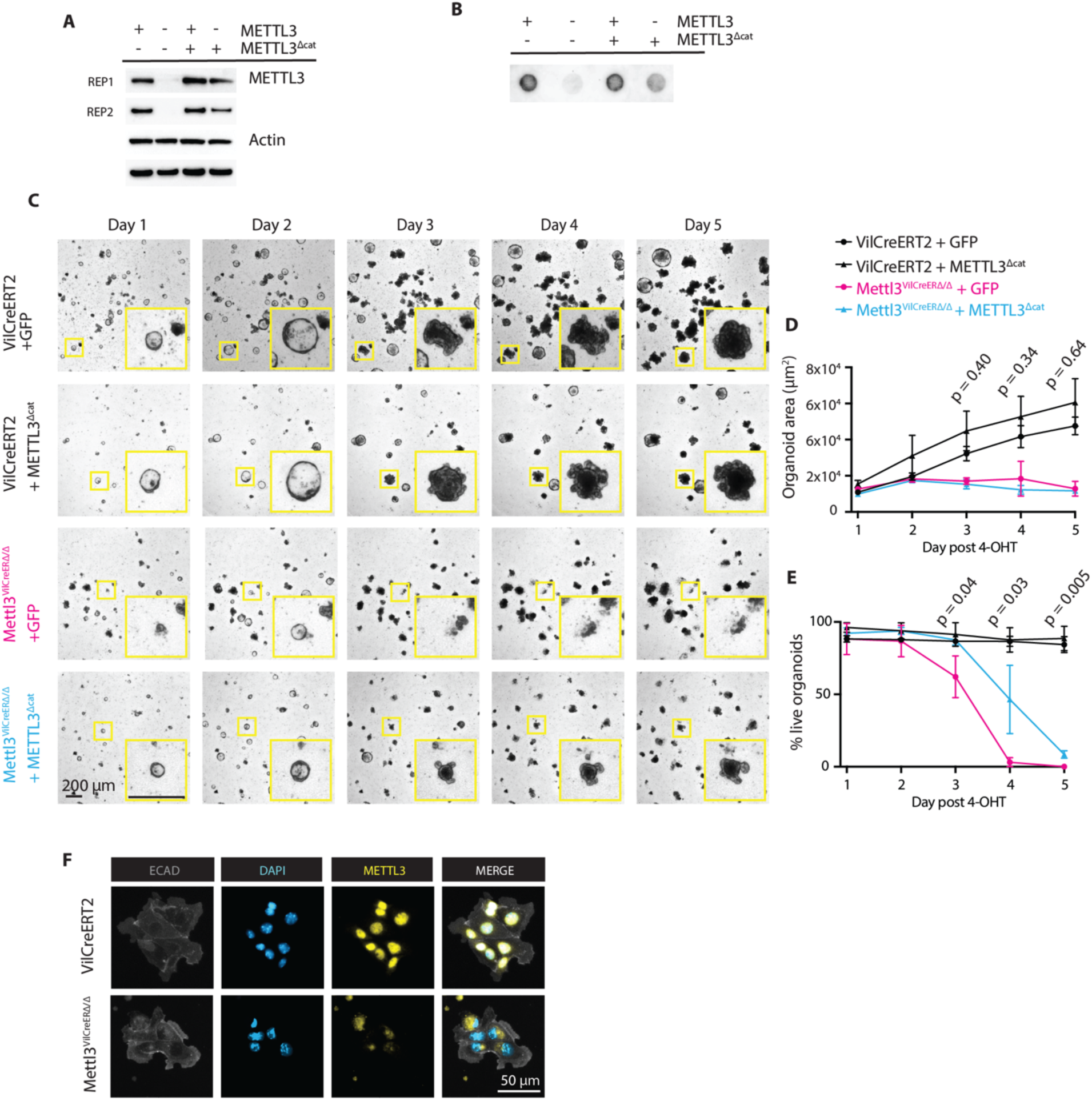
Catalytic inactive METTL3 does not rescue death of *Mettl3^VilCreERΔ/Δ^* enteroids. **(A)** Western for METTL3 two days post 4-OHT. **(B)** m^6^A dot blot in enteroids three days post 4- OHT. Each dot is 35 ng isolated mRNA. **(C)** Representative images of enteroid appearancein the five days post 4-OHT treatment. Individual enteroids are highlighted in yellow insets. **(D)** ImageJ quantification of average enteroid 2D area at each day post 4-OHT treatment from (C). Each point represents n=3 passage- separated biological replicates. Data presented as mean +/- SD. **(E)** Percent live enteroids at each day post 4-OHT treatment in enteroids from (C). Each point represents n=3 passage-separated biological replicates. Data presented as mean +/- SD. For D & E, P-value represents unpaired parametric Student’s t test on days 3, 4, 5. P-values shown are for the comparison between *Mettl3^VilCreERΔ/Δ^* + GFP vs *Mettl3^VilCreERΔ/Δ^* + METTL3^Δcat^. **(F)** Whole mount staining of METTL3 in intestinal epithelial monolayers two days post 4-OHT treatment.

**Supplemental Figure 5.**
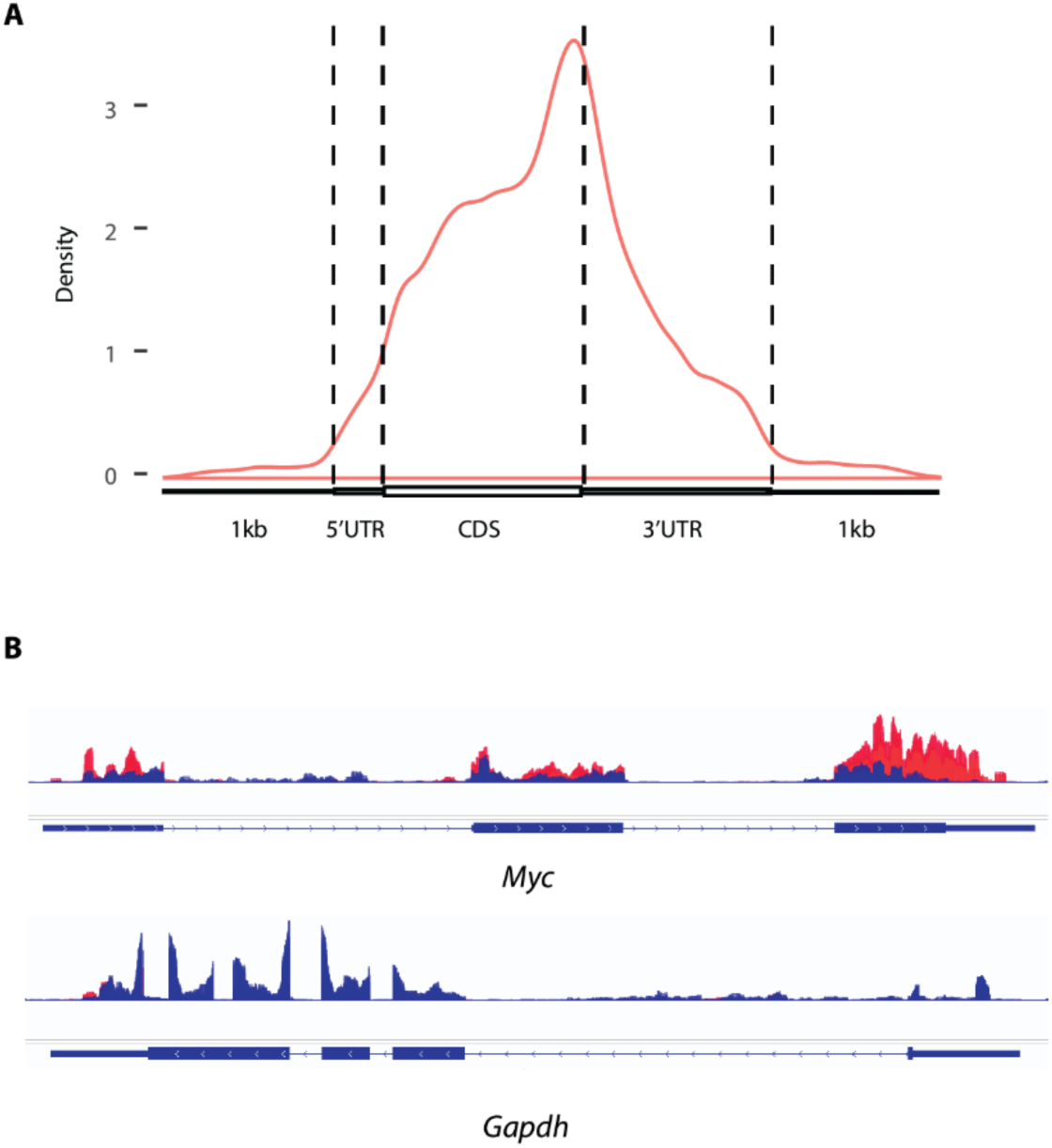
Quality control for m^6^A-seq. **(A)** Metagene density plot depicting distribution of m^6^A peaks called by Exomepeak2 within m^6^A-seq data from wildtype mouse small intestinal crypt epithelium. **(B)** m^6^A-seq read density (red) compared to input RNA read density (blue) for positive control (*Myc)* and negative control (*Gapdh*) transcripts as seen in Integrated Genomics Viewer.

**Supplemental Figure 6.**
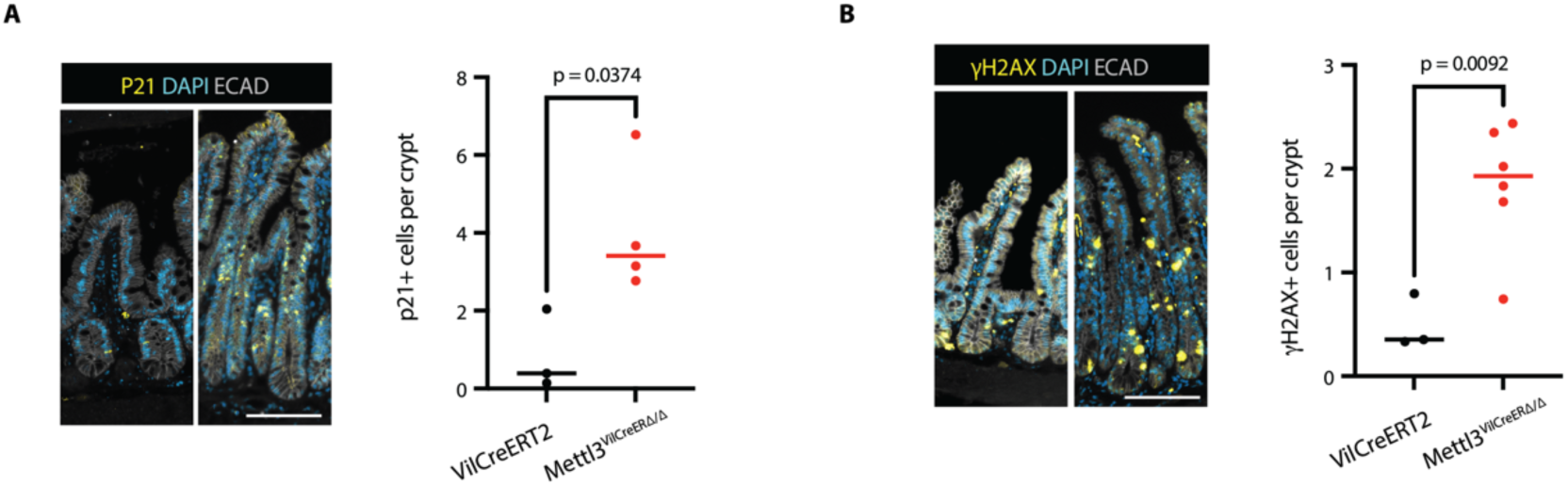
Increases in senescence and genomic instability after METTL3 deletion are significant compared to *Villin-CreERT2* controls. (A and B) Representative images and quantification of p21 (n=3,4) and γH2AX (n=3, 6) staining in distal half small intestine of *Villin-CreERT2* and *Mettl3^VilCreERΔ/Δ^*mice two days post final tamoxifen injection. Images and quantification from areas of most severe histological distortion in distal small intestine of mice two days post final tamoxifen injection. Each data point is the mean of three representative sections imaged per mouse with bar at median value and p denotes value of unpaired parametric Student’s t test. Scale bars 100 µM.

**Supplemental Table 1.** Full list of primary and secondary antibodies used in this study.

| Primary antibody- IF | Catalog number | Antigen retrieval condition (if applicable) |
| --- | --- | --- |
| Rabbit anti-METTL3 | abcam ab195352 | Tris-EDTA pH 9.0 |
| Rabbit anti-p21 | Proteintech 28248-1-AP | Tris-EDTA pH 9.0 |
| Rabbit anti-Phosphorylated-Histone H2A.X (Ser139) | CST 9718 | Tris-EDTA pH 9.0 |
| Rabbit anti-SP-1 Chromogranin A | Immunostar 20085 | Tris-EDTA pH 9.0 |
| Rabbit anti-LYZ1 | Dako A0099 | Citric Acid pH 6.0 |
| Rabbit anti-MUC2 | Genetex GTX100664 | Citric Acid pH 6.0 |
| Rabbit anti-Ki67 | abcam ab16667 | Citric Acid pH 6.0 |
| Goat anti-E-Cadherin | R&D Systems AF748 | Either antigen retrieval method |
| Mouse anti-beta-Actin | Sigma A5316 |  |
| Rabbit anti-METTL14 | Sigma HPA038002 |  |
| Rabbit anti-UBE1 | Proteintech 15912-1-AP |  |
| Rabbit anti-SEC13 | Proteintech 15397-1-AP |  |
| Mouse anti-c-Yes | Santa Cruz sc-8403 |  |
| Rabbit anti-KRAS | Proteintech 12063-1-AP |  |
| **Secondary antibody- IF** |  |  |
| Alexa Fluor 488 Anti-goat IgG | Jackson 805-545-180 |  |
| Cy3 Anti-Rabbit IgG | Jackson 711-165-152 |  |
| **Secondary antibody- Western** |  |  |
| Anti-rabbit HRP | CST 7074 |  |
| Anti-mouse HRP | Novus NBP1-75249 |  |

**Supplemental Table 2.** Histopathological scoring rubric adapted from (55) used for small intestine and colonic histopathological scoring, with scoring rules added for villus damage (villus damage only assessed in small intestine).

| Feature graded | Grade | Description |
| --- | --- | --- |
| Inflammation | 0 | None |
|  | 1 | Slight |
|  | 2 | Moderate |
|  | 3 | Severe |
| Extent | 0 | None |
|  | 1 | Mucosa |
|  | 2 | Mucosa and submucosa |
|  | 3 | Transmural |
| Regeneration | 4 | No tissue repair |
|  | 3 | Surface epithelium not intact |
|  | 2 | Regeneration with crypt depletion |
|  | 1 | Almost complete regeneration |
|  | 0 | Complete regeneration or normal tissue |
| Crypt damage | 0 | None |
|  | 1 | Basal 1/3 damaged |
|  | 2 | Basal 2/3 damaged |
|  | 3 | Only surface epithelium intact |
|  | 4 | Entire crypt and epithelium lost |
| Villus damage (SI only) | 0 | No change in villus height |
|  | 1 | 25% reduction in villus height |
|  | 2 | 50% reduction in villus height |
|  | 3 | 75% reduction in villus height |
|  | 4 | Complete loss of villus |
| Percent involvement | 1 | 1-25% |
|  | 2 | 26-50% |
|  | 3 | 51-75% |
|  | 4 | 76-100% |

**Supplemental Table 3.** Full list of Taqman qPCR primers used in this study.

| Gene | Assay number |
| --- | --- |
| *Actb* | Mm02619580_g1 |
| *Olfm4* | Mm01320260_m1 |
| *Bmi1* | Mm03053308_g1 |
| *Clu* | Mm01197002_m1 |
| *Lgr5* | Mm00438890_m1 |
| *Ascl2* | Mm01268891_g1 |
| *Cd44* | Mm01277161_m1 |
| *Hopx* | Mm00558630_m1 |
| *Kras* 3' UTR (also used for total transcript) | Mm00517494_m1 |
| *Kras* 5' UTR | Mm00517491_m1 |
| *Gapdh* | Mm99999915_g1 |

**Supplemental Data 1. (separate file). RNA-seq and Ribo-seq in METTL3 KO enteroids.** Full results of RNA-seq and Ribo-seq from n=3 *Villin-CreERT2* (CTRL) and n=3 *Mettl3^VilCreERΔ/Δ^* (KO) enteroid biological replicates 72 hours after initiating 4-OHT treatment. Green columns and blue columns display transcripts per million (TPM) values output by Kallisto in total RNA and ribosome footprint RNA fractions, respectively. Light orange columns correspond to translational efficiency (TE) values for each transcript determined by dividing the TPM in the total RNA library by the TPM in the ribosome footprint library for each individual transcript and sample. P value refers to the comparison between mean TE of CTRL vs mean TE of KO replicates.

**Supplemental Data 2. (separate file). m6A-seq in wildtype mouse crypt epithelium.** Full output of exomePeak2 analysis of m^6^A-sequencing data produced in epithelial cells sorted from distal small intestinal crypts of n=3 adult wildtype mice.

