## Supplementary material for "Intestinal transit amplifying cells require METTL3 for growth factor signaling, KRAS expression, and cell survival": BioRender License

49 Spadina Ave. Suite 200
Toronto ON M5V 2J1 Canada
www.biorender.com

### Confirmation of Publication and Licensing Rights

**April 6th, 2023**
**Science Suite Inc.**

***Subscription:*** *Institution*
***Agreement number:*** *NL257PX0BL*
***Journal name:*** *bioRxiv*

To whom this may concern,

This document is to confirm that Charles Danan has been granted a license to use the BioRender content, including icons, templates and other original artwork, appearing in the attached completed graphic pursuant to BioRender's Academic License Terms. This license permits BioRender content to be sublicensed for use in journal publications.

All rights and ownership of BioRender content are reserved by BioRender. All completed graphics must be accompanied by the following citation: "Created with BioRender.com".

BioRender content included in the completed graphic is not licensed for any commercial uses beyond publication in a journal. For any commercial use of this figure, users may, if allowed, recreate it in BioRender under an Industry BioRender Plan.

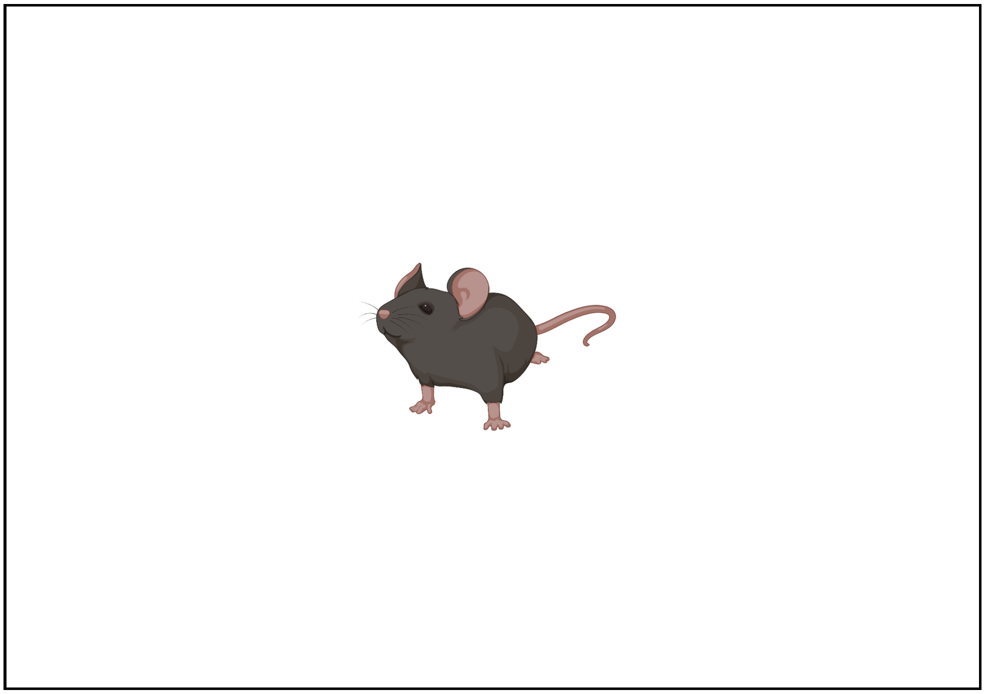

*For any questions regarding this document, or other questions about publishing with BioRender refer to our BioRender Publication Guide, or contact BioRender Support at.*
